# Interaction capacity underpins community diversity

**DOI:** 10.1101/2020.04.08.032524

**Authors:** Masayuki Ushio

**Affiliations:** Hakubi Center, Kyoto University, Kyoto 606-8501, Japan / Center for Ecological Research, Kyoto University, Otsu 520-2113, Japan

**Keywords:** Communidy diversity, Connectance, Ecosystem monitoring, Empirical dynamic modeling, Interaction capacity, Interaction strength

## Abstract

How patterns in community diversity emerge is a long-standing question in ecology. Theories and experimental studies suggested that community diversity and interspecific interactions are interdependent. However, evidence from multitaxonomic, high-diversity ecological communities is lacking because of practical challenges in characterizing speciose communities and their interactions. Here, I analyzed time-varying causal interaction networks that were reconstructed using 1197 species, DNA-based ecological time series taken from experimental rice plots and empirical dynamic modeling, and show that species interaction capacity, namely, the sum of interaction strength that a single species gives and receives, underpins community diversity. As community diversity increases, the number of interactions increases exponentially but the mean species interaction capacity of a community becomes saturated, weakening interaction among species. These patterns are explicitly modeled with simple mathematical equations, based on which I propose the “interaction capacity hypothesis,” namely, that species interaction capacity and network connectance are proximate drivers of community diversity. Furthermore, I show that total DNA copy number and temperature influence species interaction capacity and connectance nonlinearly, explaining a large proportion of diversity patterns observed in various systems. The interaction capacity hypothesis enables mechanistic explanations of community diversity, and how species interaction capacity is determined is a key question in ecology.

## Introduction

How patterns in community diversity in nature emerge is one of the most challenging and long-standing questions in ecology (Gaston 2000). Community diversity, or species diversity, a surrogate of biodiversity that is most commonly focused on (Willig *et al*. 2003), is a collective consequence of community assembly (Vellend 2016). In the community assembly processes, interspecific interactions, which contribute to the process of selection, play an important role in shaping community diversity particularly at a local (i.e., small/short spatiotemporal) scale (Kokkoris *et al*. 1999; Maynard *et al*. 2018). Therefore, they have played a central role when devising theories of ecological communities, including modern coexistence theory (Chesson 2000), niche theory (Chase & Leibold 2003) and many others. Understanding how interspecific interactions shape community diversity is key to understanding how patterns in an ecological community emerge in nature.

Theoretical studies and simple manipulative experiments have supported the view that interspecific interactions contribute to community diversity (May 1972; Kokkoris *et al*. 1999; Allesina & Tang 2012; Mougi & Kondoh 2012; Reynolds & Bruno 2013; Bairey *et al*. 2016; Ratzke *et al*. 2020). Nonlinear, state-dependent interspecific interactions have been shown to influence community diversity, composition and even dynamics (Reynolds & Bruno 2013; Bairey *et al*. 2016; Ushio *et al*. 2018a), and weak interactions are key to the maintenance of community diversity (Wootton & Emmerson 2005; Ratzke *et al*. 2020). However, despite enormous efforts to understand the interdependence between interspecific interactions and community diversity, especially efforts made in theoretical studies (May 1972; Allesina & Tang 2012; Mougi & Kondoh 2012; e.g., Ratzke *et al*. 2020), whether and how interspecific interactions control diversity in a complex, high-diversity, empirical ecological community remain poorly understood. This is largely due to two difficulties: (i) the number of species and interspecific interactions examined in previous theoretical and experimental studies have been limited compared with those of a real, high-diversity ecological community under field conditions, and (ii) detecting causal relationships between interspecific interactions and diversity is not straightforward because manipulative experiments, a most effective strategy to detect causality, are not feasible when a large number of species and interactions are targeted under field conditions. Nonetheless, understanding the mechanism by which interspecific interactions drive community diversity in nature is necessary for predicting responses of ecological communities and their functions to the ongoing global climatic and anthropogenic threats (Aronson *et al*. 2014).

To overcome the previous limitations and examine the causal relationships between interspecific interactions and diversity of a specious, empirical community, I integrated quantitative environmental DNA monitoring and a nonlinear time series analysis, and found that the capacity of species for interspecific interactions underpins community diversity. Efficient water sampling, DNA extraction and quantitative MiSeq sequencing with an internal standard DNA (Ushio *et al*. 2018b; Ushio 2019) overcame the first difficulty noted above: quantitative, highly diverse, multitaxonomic, daily, 122-day-long ecological time series were obtained from five experimental rice plots under field conditions. This extensive ecological time series was analyzed using a framework of nonlinear time series analysis, empirical dynamic modeling (EDM) (Sugihara *et al*. 2012; Ye *et al*. 2015a; Deyle *et al*. 2016), to overcome the second difficulty: EDM quantified fluctuating interaction strengths, reconstructed the time-varying interaction network of the ecological communities, and detected causal relationships between network properties and community diversity. Here, I look specifically at how interaction strengths change with community diversity, and how interspecific interactions and community diversity are causally coupled. Then, I derive a hypothesis that explains community diversity in various systems, which I call the “interaction capacity hypothesis.”

### Experimental design and ecological community monitoring

Ecological time series were taken from five experimental rice plots established at the Center for Ecological Research, Kyoto University, Japan (Fig. 1A, Fig. S1). Ecological communities were monitored by analyzing DNA in water samples taken from the rice plots using two types of filter cartridge (Ushio 2019). Daily monitoring during the rice growing season of 2017 (23 May to 22 September) resulted in 1220 water samples in total (5 plots × 2 filter types [*ϕ* 0.45-*μ*m and *ϕ* 0.22-*μ*m filters] × 122 days). Prokaryotes and eukaryotes (including fungi and animals) were analyzed by amplifying and sequencing 16S rRNA, 18S rRNA, ITS (for which DNAs extracted from *ϕ* 0.22-*μ*m filters were used), and mitochondrial COI regions (for which DNAs from*ϕ* 0.45-*μ*m filters were used), respectively, using quantitative MiSeq sequencing (Ushio *et al*. 2018b; Ushio 2019). Over 80 million reads were generated by four runs of MiSeq, and the sequences generated were then analyzed using the amplicon sequence variants (ASVs) method (Callahan *et al*. 2016) (see Fig. S2 and Supplementary Information for the quality of the sequencing). Examination of the relationships between the copy numbers and sequence reads of standard DNAs (Fig. S3A, B) and comparisons of the quantitative MiSeq method with the total DNA quantifications, quantitative PCR, and shotgun metagenomic analysis showed that the quantitative MiSeq sequencing has a reasonable capability to measure the quantity, composition, and diversity of the ecological communities (Fig. S3C-I). The ecological time series generated in this study were critically different from the relative abundance data commonly generated in microbiome studies because my data includes information on the absolute abundance. In addition, although estimations of DNA copy numbers may be biased due to experimental limitations (e.g., overestimations/underestimations of DNA copy numbers of a particular ASV due to DNA extraction and PCR biases), such biases would not be critical for subsequence time series analysis because it analyzes fluctuations in the ecological time series (i.e., time series were standardized before the time series analysis). Among over 10000 ASVs detected, 1197 ASVs (equivalent to 1197 taxonomic units) were abundant, frequently detected and contained enough temporal information for subsequent time series analyses (see Methods).

**Figure 1.**
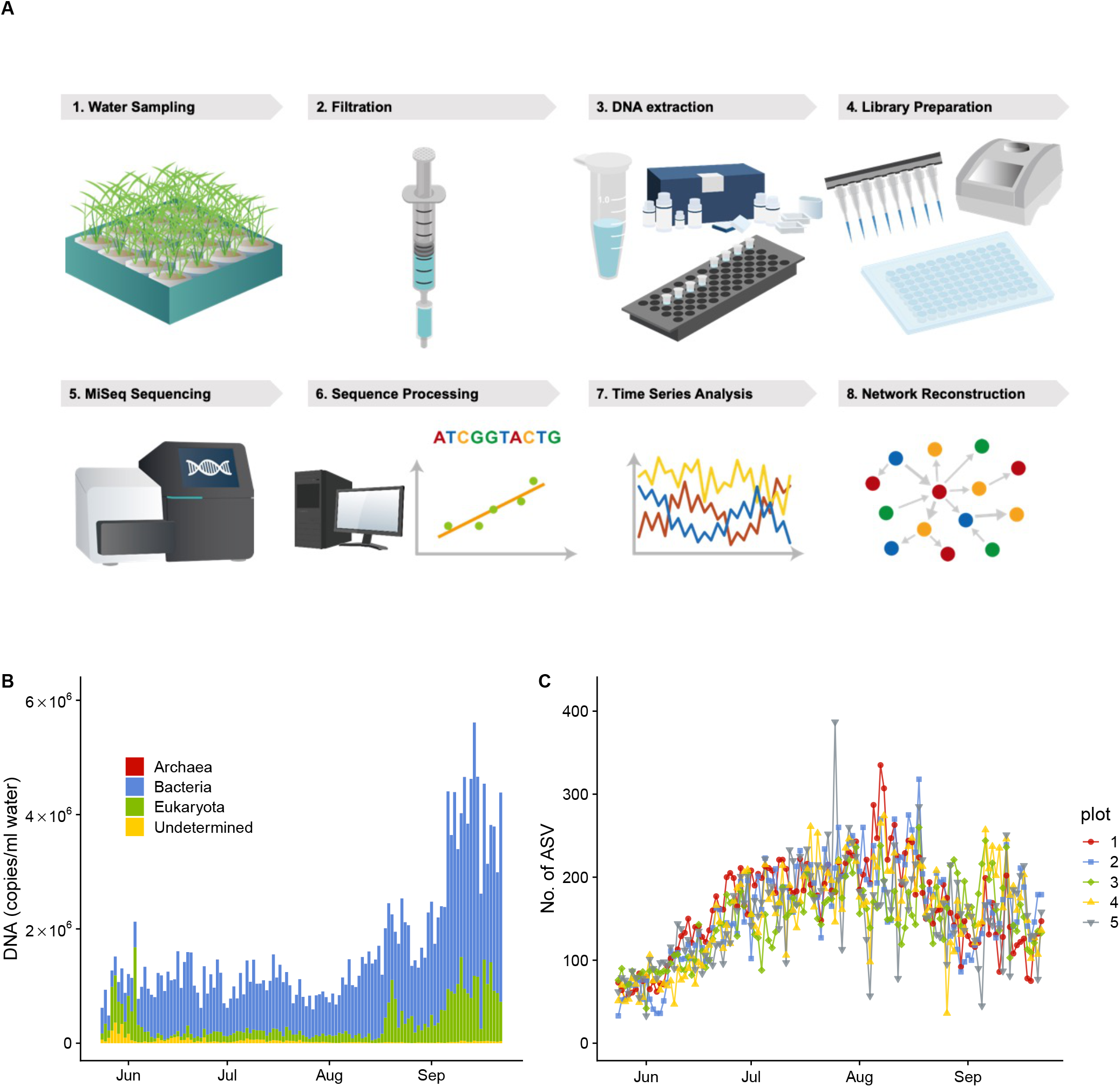
Workflow of the present study and time series of the rice plot ecological communities. **A** Workflow of the present study. **B** Mean DNA concentrations of the ecological communities in rice plots. Different colors indicate different superkingdoms. **C** Temporal patterns of the number of ASVs detected from each plot. Different symbols and colors indicate different rice plots.

### Community dynamics and reconstruction of fluctuating interaction network

The total DNA copy number increased late in the sampling period (Fig. 1B). In contrast, ASV diversity (a surrogate of species diversity in the present study) was highest in August and then decreased in September (Fig. 1C). Prokaryotes largely accounted for this pattern (Fig. S4), which is not surprising given their higher diversity and abundance compared to the other taxa.

Fluctuating interaction networks were reconstructed using EDM, a time series analytical framework for nonlinear dynamics (Sugihara *et al*. 2012; Ye *et al*. 2015a; Deyle *et al*. 2016). In the analysis, I detected causally related ASV pairs using convergent cross mapping (CCM) (Sugihara *et al*. 2012; Osada & Ushio 2021), a causality test of EDM, and then quantified the interaction strengths by multivariate, regularized S-maps (Sugihara 1994; Deyle *et al*. 2016; Cenci *et al*. 2019). Linear trends of air temperature during the monitoring season were included in the S-maps, and thus the interaction strength estimated here reflect net interactions between species (see Methods). In addition, although the total number of ASVs was over 1000, most ASVs have fewer than 20 causal interactions (Fig. S5B) suggesting that the estimation of interaction strengths by S-map would be reliable (i.e., the number of data points is more than or roughly equal to the square of the dimensions of reconstructed state space, which is required for robust estimations of the S-map coefficients).

Figure 2 shows the reconstructed network of the detected interactions over the monitoring period (for the time-varying interaction networks, see Fig. S5A and Supplementary Video). The properties of the ecological network changed over time. For example, relatively dense interactions among community members in July and August (in Plot 1) disappeared by September. Interestingly, dynamic stability (Ushio *et al*. 2018a), an index that quantifies how fast the community bounces back from small perturbations, was almost always over 1, suggesting unstable community dynamics (Fig. S5C-G). This pattern may not be surprising because the rice plots were open systems under field conditions. Many community members could immigrate and emigrate, leading to inherently unstable community dynamics. Analysis of the dynamic stability of subset communities suggested that fluctuations in moderately abundant community members could contribute to the unstable dynamics (Fig. S5D-G). The fluctuating, seemingly unstable conditions might be key to understanding the coexistence of many community members in nature. Alternatively, calculating and comparing different stability measures such as structural stability (Cenci & Saavedra 2019) may provide a different insight.

**Figure 2.**
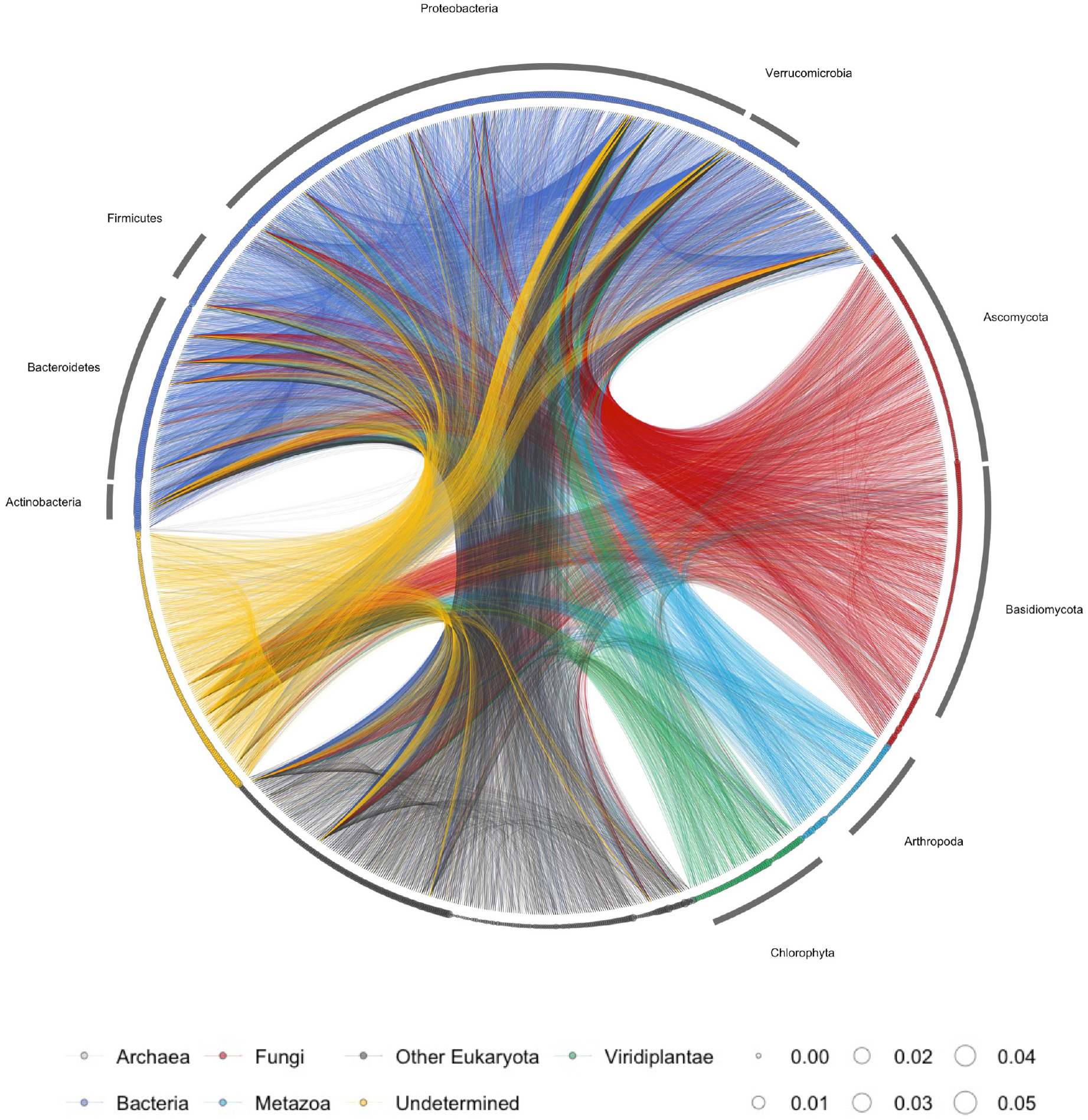
Reconstructed interaction network of the ecological community. Lines indicate causal influences between nodes, and line colors indicate causal taxa (e.g., blue lines indicate the causal influences from bacterial species to another species). The size of each node (circle) represents the DNA copy number of the ASV (n = 1197 for nodes). Different colors of nodes indicate different taxa, as shown at the bottom. Note that, although interaction strengths were quantified at each time point, the information on the time-varying interactions are not shown in the network. The detailed, daily fluctuating interaction network is presented in Fig. S5A and https://github.com/ong8181/interaction-capacity as an animation.

### Patterns emerging in the interaction networks

Properties of the interaction networks showed intriguing patterns. As ASV diversity increases, the mean interaction strength per link decreases (Fig. 3A; for mathematical definitions of network properties, see Methods), while the number of interactions in a community increases exponentially (Fig. S6), being consistent with the theoretical and experimental evidence (Kokkoris *et al*. 1999; Ratzke *et al*. 2020). This suggests that the total interaction strength that a species receives and gives might not exceed a certain upper limit, which I call “interaction capacity,” even when ASV diversity and the number of interactions in a community increase. The availability of time and energy, which are required to interact with other species, are limited. For example, it seems difficult for a single species to strongly interact with a large number of species within a certain time interval if their body size and generation time are in a similar range. Also, interspecific interactions usually involve consumption of a certain amount of energy. Thus, it is intuitively plausible to assume that there is a certain upper limit of interaction capacity which is the product of the interaction strength and the number of interactions. Indeed, the upward trend of the mean interaction capacity is weakened when ASV diversity is over 100 in the studied system (Fig. 3B), supporting the assumption that there is an upper limit for the interaction capacity.

**Figure 3.**
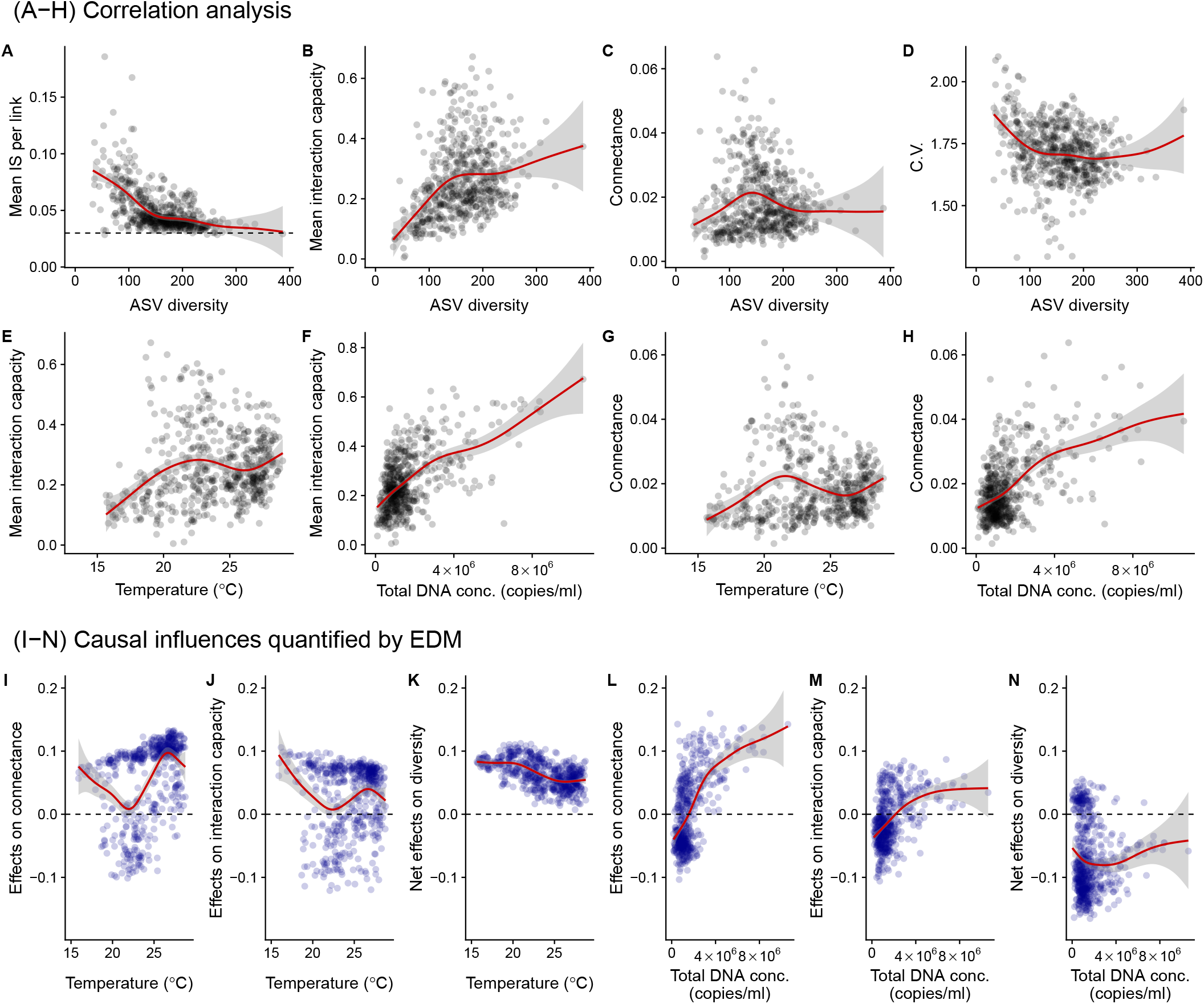
Relationships between the interaction network properties. **A**-**D** Co-varying relationships (correlations) between ASV diversity and properties of the interaction network, i.e., the mean interaction strength (**A**), interaction capacity (**B**), connectance (**C**) and coefficients of variations in population dynamics (**D**). Interaction capacity is defined as the sum of absolute values of interaction strength that a species gives or receives. Dashed line in A indicates a converged value of mean interaction strength (≈ 0.03). **E**-**H** Relationships between interaction capacity, connectance, mean air temperature and total DNA copy numbers. **I**-**N** Causal influences of air temperature and total DNA copy numbers on connectance, interaction capacity and community diversity quantified by empirical dynamic modeling (EDM). Convergent cross mapping (CCM) was first applied to each pair, and then multivariate, regularized S-map was applied to quantify the causal influences. Red lines indicate statistically clear nonlinear regressions by general additive model (GAM) and gray shaded region indicate 95% confidence interval.

Another important property of the interaction networks, connectance, is also relatively constant as ASV diversity varies (Fig. 3C; for the definition of connectance, see Methods). Importantly, these patterns were not reproduced when the randomly shuffled version of the original time series was analyzed (Fig. S7). In addition, this pattern of interaction strength per link decreasing and converging when the number of interactions and/or species is high is valid even at the species level (Fig. S8A). These findings suggest that the original results may not have been experimental or statistical artifacts but rather may have emerged as consequences of empirical community assembly processes. Another intriguing pattern is that mean values of coefficients of variation (C.V.) of DNA copy numbers, an EDM-independent index of realized temporal variability, decreases as a function of ASV diversity (Fig. 3D; one outlier shows a relatively high C.V. regardless of a high community diversity). This showed, for the first time, that the small temporal variability in species abundances observed in diverse plant communities (Tilman *et al*. 2006) is also valid even when all major cellular organisms are taken into account, and that there is a connection between community diversity and temporal dynamics in this system.

### Direct coupling of community diversity and interaction capacity revealed by simple mathematical equations

To better understand and clarify the implications of the patterns that emerged in the network properties, I explicitly show the relationship between the network properties by developing a simple mathematical model. By starting with minimal assumptions, I demonstrate that community diversity and interaction capacity are interdependent on each other. Since connectance, *C*, is defined as *C* = *N*_*link*_ / *S*^2^, species diversity (species richness), *S*, can be simply represented as 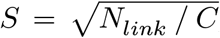, where *N*_*link*_ indicates the total number of interactions in a community. Furthermore, *N*_*link*_ can be decomposed into the mean interaction capacity at the community level (*IC*), defined as 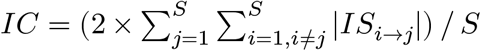 (see Methods), and the mean interaction strength per link, *IS*_*link*_, as follows:

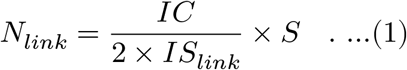

*IC*/*IS*_*link*_ is divided by 2 because each interaction strength is counted twice (for donor and receiver species). Therefore, *S, C, IC* and *IS*_*link*_ should satisfy the following relationship:

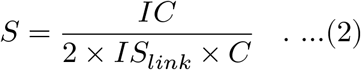

No system-specific assumption is used to derive Eqn. (2), and thus *S, IC, IS*_*link*_ and *C* should satisfy Eqn. (2) in any system and under any condition. Note that the four parameters are all interdependent, and thus a change in one parameter influences other parameters. Causal tests using the time series taken from the rice plots suggested that the four parameters in Eqn. [2] are indeed causally interdependent in the empirical communities (Fig. S9).

The simple equation, Eqn. [2], suggests that there is a negative relationship between *S* and *C* or *IS*_*link*_ given that the other parameters are constant. Similar arguments were made in previous seminal theoretical studies (May 1972; Allesina & Tang 2012), but there are important contributions of Eqn. [2] to understanding empirical ecological communities. “Interaction capacity” is an easily understandable concept, and it can be defined as “trait” for any biological level (e.g., for community, species, or even local populations). Eqn. [2] explicitly shows the linkage between the interaction capacity and other fundamental properties of an ecological community, opening up a new research direction to link the biological trait and network properties. Furthermore, it enables intuitive and clear explanations of mechanisms of community diversity by investigating how *IC* and *C* are determined and it has empirical supports as shown in the following sections.

### The interaction capacity hypothesis

A potentially important feature of the emerging empirical patterns is that *IS*_*link*_ tends to converge when community diversity increases (Fig. 3A). When using the system-specific parameter value of converged *IS*_*link*_ in the present study, maximum possible community diversity in the rice plots is approximated based on Eqn. [2] as follows:

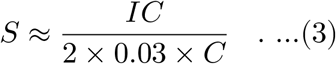

Eqn. [3] predicts that the interaction capacity should have positive influences on community diversity. Further, connectance (or mean interaction strength) should have negative influences on community diversity, which is consistent with previous theoretical studies (e.g., May 1972; Allesina & Tang 2012). Therefore, this mathematical model indicates that species diversity can be predicted if we know how *IC* and *C* are determined in a community.

My ecological time series provides a unique opportunity to elucidate how interaction capacity (*IC*) and connectance (*C*) are determined under field conditions. In the analysis, I focused on two fundamental variables that are statistically independent of the network properties: air temperature and total DNA copy number (an index of total abundance/biomass). Also, I focused on these because air temperature is an independent external driver of community dynamics and because total abundance could be an index of net ecosystem productivity or available energy in a system, which could be a potential driver of community diversity (Evans *et al*. 2005; Huston 2014). Correlation analysis shows that the interaction capacity is positively correlated with mean air temperature and total DNA copy number (Fig. 3E,F). Connectance is positively correlated with total DNA copy number, and is weakly correlated with mean air temperature (Fig. 3G,H).

Causal relationships between the network properties and external forces were examined with CCM, and the results suggested that mean interaction capacity and connectance are causally influenced by mean air temperature and total DNA copy number (Fig. S9). The S-map revealed that mean air temperature in general positively influenced interaction capacity and connectance, as indicated by mostly positive values along the gradient (Fig. 3I,J; values on the y-axis indicate how changes in temperature cause changes in interaction capacity or connectance). Temperature may influence many aspects of biological processes, e.g., physiological rates of individuals, and therefore, the influences of air temperature on interaction capacity and connectance may arise from the increased activities of individuals. Although temperature effects on connectance and mean interaction capacity at the community level were comparable, the net effects of temperature on diversity, that is, effects of temperature through its effects on interaction capacity and connectance, were consistently positive (Fig. 3K). This indicates that the positive influence of temperature on mean interaction capacity is stronger than that on connectance for shaping community diversity.

When total DNA copy number is low, its influence on connectance is variable (Fig. 3L). When total DNA copy number is high, however, total DNA copy number strongly and positively influences connectance, suggesting that denser populations should have higher connectance, probably because the greater population size may facilitate random encounters among individuals or species. On the other hand, the influences on interaction capacity are relatively small and variable (Fig. 3M). These results predict that total DNA copy number (or abundance/biomass) over a certain threshold negatively influence diversity, which is indeed the case here (Fig. 3N).

Together, the results of EDM show that interaction capacity and connectance are influenced by temperature (*T*) and total DNA concentration (*DNA*), suggesting that species diversity, *S*, can be approximated using these two fundamental parameters as follows:

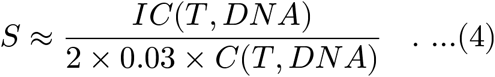

Although the influences of temperature and abundance (biomass or energy) on diversity have long been recognized in the literatures (e.g, Begon *et al*. 2005), this model, supported by empirical evidence, provides mechanistic explanations about how temperature and abundance control community diversity. The present study did not include other potentially important abiotic factors such as water pH and nutrient availability, but the mechanisms of the influence of such factors may also be understood by considering their effects on interaction capacity and connectance. Because community diversity, interaction capacity and connectance are interdependent in any system and under any condition according to Eqn. [2], the influences of any biotic/abiotic factors on community diversity can be mechanistically explained and predicted if we can understand the influences of these factors on interaction capacity and connectance, a proposal which I call the “interaction capacity hypothesis.”

### Empirical evidence supporting the interaction capacity hypothesis in other systems

The simple mathematical model and the analyses of the extensive ecological time series reported here suggest that proximate drivers of community diversity are interaction capacity and connectance, and that the long-recognized patterns that temperature and total species abundance influence community diversity are underpinned by their influences on interaction capacity and connectance. In some cases, variable responses of community diversity to temperature and/or abundance might be observed because of the nonlinear influences of temperature and abundance on interaction capacity and connectance (Fig. 3I-N). Conversely, if the present results can be generalized to other systems, it should be possible to explain community diversity reasonably well by a nonlinear regression using temperature and total abundance.

To verify this expectation, I compiled published data from various ecosystems. The collected data include two global datasets and four local datasets collected in Japan: (i) global ocean microbes (Sunagawa *et al*. 2015), (ii) global soil microbes (Bahram *et al*. 2018), (iii) fish from a coastal ecosystem (Masuda 2008), (iv) prokaryotes from freshwater lake ecosystems (Okazaki *et al*. 2017), (v) zooplankton from a freshwater lake ecosystem (Sakamoto *et al*. 2018) and (vi) benthic macroinvertebrates from freshwater tributary lagoon ecosystems (Okano *et al*. 2018). Because the influences of temperature and total species abundance/biomass on community diversity (or interaction capacity and connectance) are likely to be nonlinear, I adopted a general additive model (Wood 2004) as follows: *log*(*S*) *s*(*log*(*T*)) + *s*(*log*(*A*)), where *S, T, A* and *s*() indicate species diversity (or OTU diversity), temperature, an index of total species abundance (or biomass) and a smoothing term, respectively. The simple, nonlinear regression using temperature and abundance (or biomass) were separately applied to each data set, which explained biodiversity surprisingly well for the five aquatic data sets (Fig. 4; adjusted *R*^2^ = 0.453-0.792) and somewhat worse for global soil data (Fig. 4B; adjusted *R*^2^ = 0.16). Connectance may be difficult to predict from abundance in spatially heterogeneous ecosystems such as soil, which may account for the lower predictability of the regression model of the soil data. Also, interaction capacity may be influenced by species identity (e.g., Fig. S8B), and in that case interaction capacity may not be simply a function of temperature and/or abundance. Thus, the accuracy with which diversity can be predicted from temperature and abundance may differ for target organisms and ecosystems. Nonetheless, biodiversity is surprisingly well explained only by a nonlinear regression using temperature and abundance, suggesting that the interaction capacity hypothesis might be applicable to a wide range of taxa and ecosystems.

**Figure 4.**
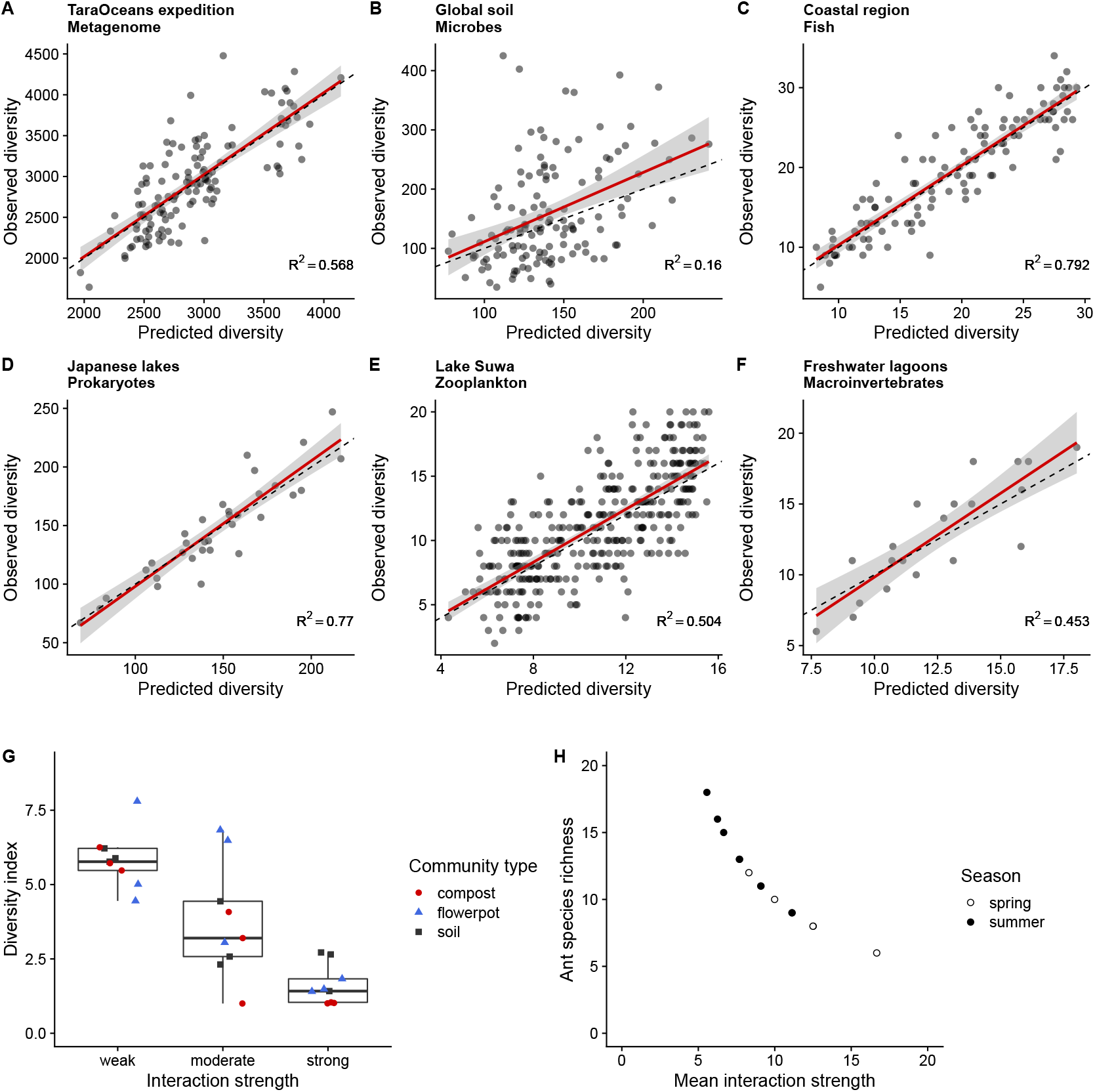
Predicted and observed diversity based on the interaction capacity hypothesis. **A**-**F** Predicted and observed diversity for metagenomic data generated by Tara Oceans research expedition (**A**), global soil microbial community (**B**), a fish community in a marine costal region in Kyoto, Japan (**C**), prokaryotic communities in Japanese lakes (**D**), zooplankton communities in Lake Suwa, Japan (**E**) and macroinvertebrate communities in freshwater tributary lagoons surrounding Lake Biwa, Japan (**F**). Values at the bottom right in panels indicate adjusted *R*^2^ of GAM. Gray shaded region indicates 95% confidence interval of the linear regression between predicted and observed diversities. **G**-**H** The negative relationship between interaction strength and community diversity. Examples from experimental microbial communities (**G**, Ratzke *et al*. 2020) and ant-plant interactions under field conditions (**H**, Yamawo *et al*. 2021).

In addition, while one piece of evidence supporting the interaction capacity hypothesis is the reasonably accurate predictions of community diversity by nonlinear regression using temperature and abundance, another supporting piece of evidence is the negative relationship between interaction strength and community diversity. To further support the interaction capacity hypothesis, I introduce two empirical results that show the negative relationship between interaction strength and community diversity (Fig. 4G, H). Ratzke *et al*. (Ratzke *et al*. 2020) manipulated the interaction strength of microbial community members by modifying medium conditions, and found that the community diversity decreased with an increase of the interaction strength (Fig. 4G). Yamawo *et al*. (Yamawo *et al*. 2021) investigated the interaction strength between ant species and a pioneer tree species, and found that increasing the local ant species diversity decreased the ant-plant interaction strength (Fig. 4H). An assumption that a species has a certain interaction capacity is one of the possible explanations for the empirical observations. Nonetheless, evidence presented here are still necessary conditions of the interaction capacity hypothesis, and further testing using independent methods and data is required.

The interaction capacity hypothesis provides quantitative and unique predictions about community diversity in nature. For example, everything else being equal, community diversity may increase with increasing temperature because of increased interaction capacity under warmer conditions (Fig. 5A, B). Similarly, community diversity will increase with increasing habitat heterogeneity because of decreased connectance (Fig. 5A, B). Thus, high diversity communities will exist under optimal environmental conditions (i.e., high interaction capacity) with spatially heterogeneous habitats (i.e., low connectance), such as those in tropical forests or soils with neutral pH (Fig. 5C, D). Furthermore, under such conditions, community dynamics will be stabilized because of the decreased interaction strengths (Figs. 3D, 5C; also see evidence from a recent experimental study Ratzke *et al*. 2020). On the other hand, low-diversity communities will exist under extreme environmental conditions with spatially homogeneous habitats, such as deserts, because of decreased (or consumed) interaction capacity and increased connectance.

**Figure 5.**
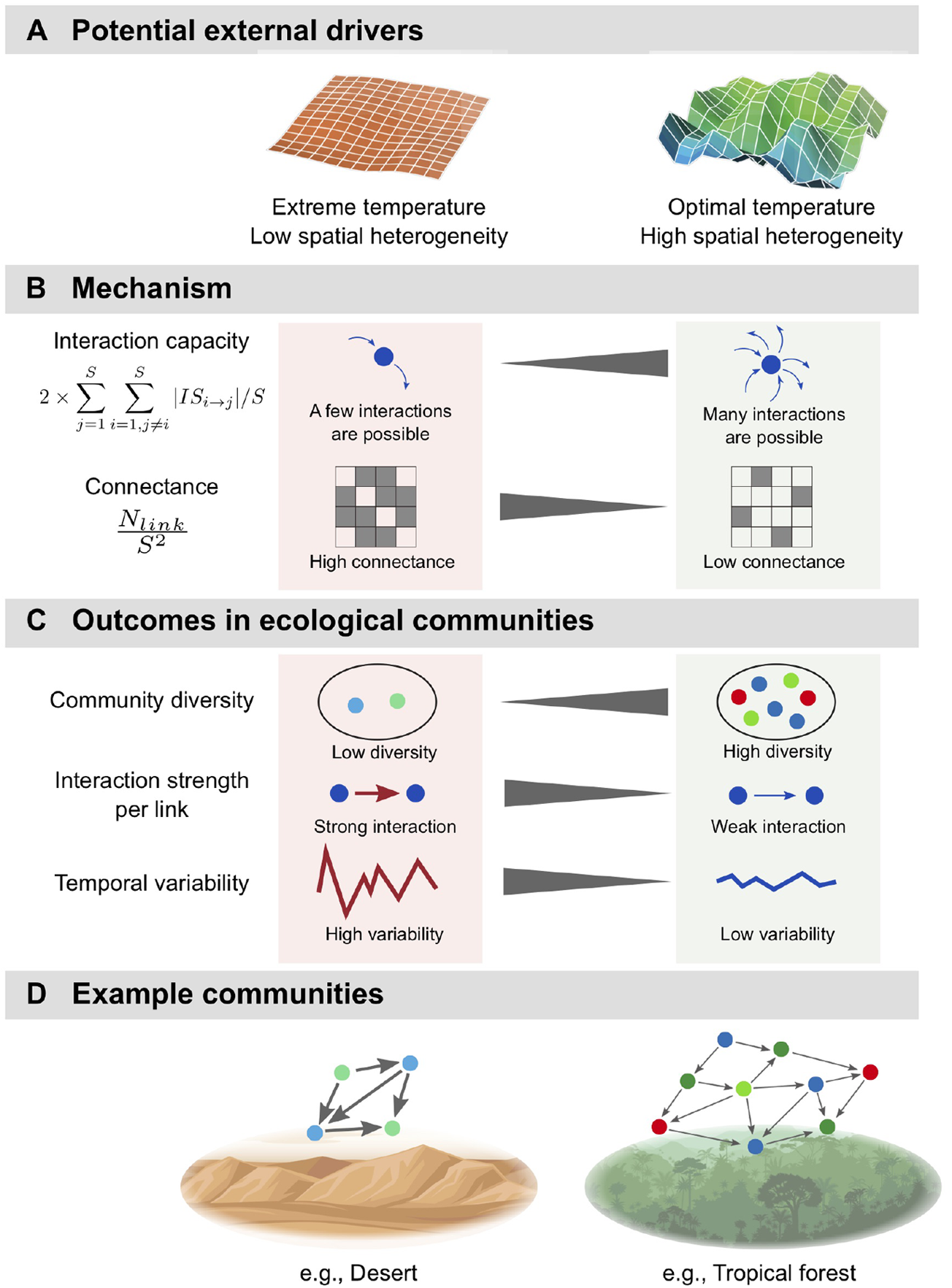
Potential drivers of the community network according to the interaction capacity hypothesis and examples of ecological communities. **A** Potential external drivers that contribute to the community diversity and network structure. **B** Mechanisms of community assembly. Extreme and optimal temperature would generally decrease and increase species interaction capacity, respectively. Spatial heterogeneity decreases connectance, which subsequently increases community diversity. **C** Outcomes of community diversity and dynamics. Extreme temperature and low spatial heterogeneity (e.g., desert ecosystem) generate a community with low diversity, high interaction strength (and a small number of interactions) and unstable dynamics. On the other hand, optimal temperature and high spatial heterogeneity (e.g., tropical forests) generate a community with high diversity, low interaction strength (but a large number of interactions) and stable community dynamics. **D** Examples of ecological communities (low-diversity versus high-diversity community) according to the potential external drivers and the interaction capacity hypothesis..

### Potential limitations of the present study

The interaction capacity hypothesis proposed in the present study seems to have reasonable supports according to the field monitoring, quantitative MiSeq sequencing, EDM, simple mathematical considerations, and meta-analysis. Nonetheless, there may be several potential limitations and I explicitly state them in this section.

First, the number of community types examined in the present study is still small. In particular, comprehensive, quantitative, and multispecies time series were taken from only one freshwater system (the artificial rice plots). Ecological communities with different community members, interaction types, and external driving forces may generate different patterns in community dynamics and interactions. For example, vertebrate species are virtually absent in my system. Because their life history and generation time are substantially different from those of species which were examined in the present study, and because detecting and quantifying interactions between organisms with different body size and time scales may be challenging, including such species may change the pattern. Interaction types (mutualism, competition, and so on) and its diversity (i.e., the ratio of positive and negative interactions in a community) might also matter (Mougi & Kondoh 2012). Differences in external driving forces (e.g., temperature and disturbance) may be another important factor that should be incorporated. Though potential outcomes of the hypothesis were confirmed by several previous studies, further studies, especially empirical ones, are needed to verify the hypothesis.

Second, there are potential experimental limitations in the present study. Because the detections of species relied on DNA sequencing, an implicit assumption of the present study is that the DNA copy number corresponds to each species abundance (note that the conversion from the DNA copy number to species abundance may be species-specific due to differences in amplification efficiencies and/or the copy number of 16S rRNA genes). Although this assumption would be reasonable for microbial species, how (environmental) DNA concentrations are related to the abundance/biomass of macro-organisms is still actively debated. Some studies demonstrated a linear relationship between DNA concentrations and species abundance for some macro-organisms (e.g., Takahara *et al*. 2012; Thomsen *et al*. 2012), but how general the positive linear relationshp is still unclear. Another potential experimental limitation of the present study is the interval of the time series. As the sampling interval was daily, interactions occurring on much shorter than the daily time scale seems to be elusive in the present study. An automated field sampling system may solve this issue (Yamahara *et al*. 2019; Hansen *et al*. 2020), but at present, such as system is not easily affordable for many field ecologists.

Lastly, there are potential statistical limitations in the present study. The number of data points influences the prediction accuracy of EDM especially when the number of variables included in the S-map becomes large (i.e., the curse of dimensionality). Although this would not cause serious problems in the present study as the number of data points (= 610) is sufficiently large compared to the dimensionality (= optimal E and the number of causal species was smaller than 14 for most species; Figs. S5B, S8A), it is still challenging to reconstruct much higher dimensional interaction networks with fewer data points (Chang *et al*. 2021). We have recently addressed this issue and developments of such statistical tools will help to understand natural, high dimensional ecosystem dynamics (Chang *et al*. 2021). Another important issue is how we can experimentally validate the results of EDM in a natural ecosystem. As described in previous studies (e.g., Sugihara *et al*. 2012), EDM was initially developed for near future predictions and causality detections in a complex, natural system where manipulative experiments are not feasible. However, building an experimental framework to validate results of EDM would be beneficial especially when a target system has a moderate size and the primary object of the analysis is the management and control of the system.

### Future directions

Considering the interdependence between interaction capacity and community diversity, how interaction capacity is determined will be a central question in ecology. For example, interaction capacity may be influenced by energy and resources provided to a system, but it can also be influenced by species identity (i.e., species’ ecology, physiology and evolutional history; e.g., higher interaction capacities of prokaryotes than of eukaryotes; see Fig. S8B). Also, rapid evolution and eco-evolutionary feedbacks may cause changes in species trait, their interactions, and dynamics (e.g., Kasada *et al*. 2014), which would influence how species assign their interaction capacity to biotic/abiotic interactions. Incorporating species identity, evolutionary history, and eco-evolutionary feedbacks into the interaction capacity hypothesis would be an interesting direction for future studies.

In addition, abiotic factors can easily and explicitly be incorporated into the interaction capacity hypothesis. For example, if interaction capacity (or energy resources) is “consumed” to adapt to harsh environmental conditions, interaction capacity that can be used for interspecific interaction will decrease, which will consequently decrease community diversity. Lastly, applying the interaction capacity hypothesis to other systems such as gene expression, neural networks and even human social networks should lead to fresh insights.

## Conclusions

How patterns in community diversity emerge has been extensively studied by experimental and theoretical approaches, yet rarely examined for highly diverse, complex ecological communities. Using DNA-based, highly frequent, quantitative, extensive ecological time series and EDM, I present evidence that interaction capacity is a key to understanding and predicting community diversity. Connectance may also play an important role, because it determines how interaction capacity is divided into each interaction link. Interaction capacity is influenced by the total abundance and temperature, which can provide mechanistic explanations for many observed ecological patterns in nature. Expanding spatial and temporal scales and incorporating the other processes in community assembly (that is, speciation, dispersal and drift) (Vellend 2016) into the interaction capacity hypothesis will further deepen our understanding of community assembly processes, which will contribute to how we can predict, manage and conserve biodiversity and the resultant ecosystem functions in nature.

## Methods

### Experimental setting

Five artificial rice plots were established using small plastic containers (90 × 90 × 34.5 cm; 216 L total volume; Risu Kogyo, Kagamigahara, Japan) in an experimental field at the Center for Ecological Research, Kyoto University, in Otsu, Japan (34° 58’ 18’ ‘ N, 135° 57’ 33’ ‘ E). Sixteen Wagner pots (*ϕ* 174.6 × *ϕ* 160.4 × 197.5 mm; AsOne, Osaka, Japan) were filled with commercial soil, and three rice seedlings (var. Hinohikari) were planted in each pot on 23 May 2017 and then harvested on 22 September 2017 (122 days; Table S1). The rice growth data are being analyzed for different purposes and thus are not shown in this report. The containers (hereafter, “plots”) were filled with well water, and the ecological community was monitored by analyzing DNA in the well water (see following subsections).

### Field monitoring of the ecological community

To monitor the ecological community, water samples were collected daily from the five rice plots. Approximately 200 ml of water in each rice plot was collected from each of the four corners of the plot using a 500-ml plastic bottle and taken to the laboratory within 30 minutes. Water samples were kept at 4°C during transport. The water was filtered using Sterivex filter cartridges (Merck Millipore, Darmstadt, Germany). Two types of filter cartridges were used to filter water samples: to detect microorganisms, *ϕ* 0.22-*μ*m Sterivex (SVGV010RS) filter cartridges that included zirconia beads inside (for degradation of the microbial cell wall) were used (Ushio 2019), and to detect macroorganisms,*ϕ* 0.45-*μ*m Sterivex (SVHV010RS) filter cartridges were used. Water in each plastic bottle was thoroughly mixed before filtration, and 30 ml and 100 ml aliquots of the water were filtered using *ϕ* 0.22-*μ*m and *ϕ* 0.45-*μ*m Sterivex, respectively (slightly adjusted when the filters were clogged). After filtration, 2 ml of RNAlater solution (ThermoFisher Scientific, Waltham, Massachusetts, USA) were added to each filter cartridge to prevent DNA degradation during storage. In total, 1220 water samples (122 days × 2 filter types × 5 plots) were collected during the census term. In addition, 30 field-level negative controls, 32 PCR-level negative controls with or without the internal standard DNAs and 10 positive controls to monitor the potential DNA cross-contamination and degradation during the sample storage, transport, DNA extraction and library preparations were used. Visual inspections of the negative and positive control results indicated no serious DNA contaminations or degradation during analyses (Fig. S2). Detailed information on the negative/positive controls are provided in Supplementary Information.

### DNA extractions

DNA was extracted using a DNeasy Blood & Tissue kit following a protocol described in my previous report (Ushio 2019). First, the 2 ml of RNAlater solution in each filter cartridge were removed from the outlet under vacuum using the QIAvac system (Qiagen, Hilden, Germany), followed by a further wash using 1 ml of MilliQ water. The MilliQ water was also removed from the outlet using the QIAvac. Then, Proteinase K solution (20 *μ*l), PBS (220 *μ*l) and buffer AL (200 *μ*l) were mixed, and 440 *μ*l of the mixture was added to each filter cartridge. The materials on the cartridge filters were subjected to cell lysis by incubating the filters on a rotary shaker (15 rpm; DNA oven HI380R, Kurabo, Osaka, Japan) at 56°C for 10 min. After cell lysis, filter cartridges were vigorously shaken (with zirconia beads inside the filter cartridges for *ϕ* 0.22-*μ*m cartridge filters) for 180 sec (3200 rpm; VM-96A, AS ONE, Osaka, Japan). The bead-beating process was omitted for *ϕ* 0.45-*μ*m cartridge filters. The incubated and lysed mixture was transferred into a new 2-ml tube from the inlet (not the outlet) of the filter cartridge by centrifugation (3500 g for 1 min). Zirconia beads were removed by collecting the supernatant of the incubated mixture after the centrifugation. The collected DNA was purified using a DNeasy Blood & Tissue kit following the manufacturer’s protocol. After the purification, DNA was eluted using 100 *μ*l of the supplied elution buffer. Eluted DNA samples were stored at −20°C until further processing.

### Library preparation

Prior to the library preparation, work spaces and equipment were sterilized. Filtered pipet tips were used, and pre-PCR and post-PCR samples were separated to safeguard against cross-contamination. PCR-level negative controls (i.e., with and without internal standard DNAs) were employed for each MiSeq run to monitor contamination during the experiments.

Details of the library preparation process are described in Supplementary Information. Briefly, the first-round PCR (first PCR) was carried out with the internal standard DNAs to amplify metabarcoding regions of prokaryotes (515F-806R primers) (Bates *et al*. 2011), eukaryotes (Euk_1391f and EukBr primers) (Amaral-Zettler *et al*. 2009), fungi (ITS1-F-KYO1 and ITS2-KYO2) (Toju *et al*. 2012) and animals (mostly invertebrates in the present study) (mlCOIintF and HCO2198 primers) (Folmer *et al*. 1994; Leray *et al*. 2013). After the purifications of the triplicate 1st PCR products, the second-round PCR (second PCR) was carried out to append indices for different templates (samples) for massively parallel sequencing with MiSeq. Twenty microliters of the indexed second PCR products were mixed, the combined library was purified, and target-sized DNA of the purified library was excised and quantified. The double-stranded DNA concentration of the library was then adjusted using MilliQ water and the DNA was applied to the MiSeq (Illumina, San Diego, CA, USA).

### Sequence processing: Amplicon sequence variant (ASV) approach

The raw MiSeq data were converted into FASTQ files using the bcl2fastq program provided by Illumina (bcl2fastq v2.18). The FASTQ files were then demultiplexed using the command implemented in Claident (http://www.claident.org) (Tanabe & Toju 2013). I adopted this process rather than using FASTQ files demultiplexed by the Illumina MiSeq default program in order to remove sequences whose 8-mer index positions included nucleotides with low quality scores (i.e., Q-score < 30).

Demultiplexed FASTQ files were analyzed using the Amplicon Sequence Variant (ASV) method implemented in the DADA2 (v1.11.5) (Callahan *et al*. 2016) package of R. First, the primers were removed using the external software cutadapt v2.6 (Martin 2011). Next, sequences were filtered for quality using the DADA2::filterAndTrim() function, and rates were learned using DADA2::learnErrors() function (MAX_CONSIST option was set as 20). Then, sequences were dereplicated, error-corrected, and merged to produce an ASV-sample matrix. Chimeric sequences were removed using the DADA2::removeBimeraDenove() function.

Taxonomic identification was performed for ASVs inferred using DADA2 based on the query-centric auto-k-nearest-neighbor (QCauto) method (Tanabe & Toju 2013) and subsequent taxonomic assignment with the lowest common ancestor algorithm (Huson *et al*. 2007) using “overall_class” and “overall_genus” database and clidentseq, classigntax and clmergeassign commands implemented in Claident v0.2.2019.05.10. I chose this approach because the QCauto method assigns taxa in a more conservative way (i.e., low possibility of false taxa assignment) than other methods. Because the QCauto method requires at least two sequences from a single microbial taxon, only internal standard DNAs were separately identified using BLAST (Camacho *et al*. 2009).

After the taxa assignment, sequence performance was carefully examined using rarefaction curves, detected reads from PCR and field negative controls (Fig. S2 and Supplementary Information). In addition, whether the PCR-based assessments of community diversity were biased was tested by performing shotgun metagenomic analysis of a few representative samples (Fig. S3F-I). Although the lower DNA concentrations and shallow sequencing depth might reduce the detection rate of rare taxa by the quantitative MiSeq sequencing, saturated rarefaction curves (Fig. S2A-D) and the results of shotgun metagenome analysis (Fig. S3F-I) suggested that the quantitative MiSeq sequencing captured most of the diversity in the water samples. Also, the results of PCR and field negative controls suggested that there were low levels of contamination during the monitoring, DNA extractions, library preparations and sequencing (Fig. S2).

### Estimations of DNA copy numbers

For all analyses in this subsection, the free statistical environment R 3.6.1 was used (R Core Team 2019). The procedure used to estimate DNA copy numbers consisted of two parts, following previous studies (Ushio *et al*. 2018b; Ushio 2019): (i) linear regression analysis to examine the relationship between sequence reads and the copy numbers of the internal standard DNAs for each sample (Fig. S3A, B), and (ii) the conversion of sequence reads of non-standard DNAs to estimate the copy numbers using the result of the linear regression for each sample. Linear regressions were used to examine how many sequence reads were generated from one DNA copy through the library preparation process. Note that a linear regression analysis between sequence reads and standard DNAs was performed for each sample and the intercept was set as zero. The regression equation was: MiSeq sequence reads = regression slope × the number of standard DNA copies [/*μ*l]. Most samples show highly significant linear relationship between the copy numbers and sequence reads of the standard DNAs (Fig. S3A, B), suggesting that the number of sequence reads produced is proportional to the copy number of DNAs within a single sample.

The sequence reads of non-standard DNAs were converted to copy numbers using sample-specific regression slopes estimated using the above regression analysis. The number of non-standard DNA copies was estimated by dividing the number of MiSeq sequence reads by the value of a sample-specific regression slope (i.e., the number of DNA copies = MiSeq sequence reads/regression slope). A previous study demonstrated that these procedures provide a reasonable estimate of DNA copy numbers using high-throughput sequencing (Ushio *et al*. 2018b).

After the conversion to DNA copy number, ASVs with low DNA copy numbers were excluded because their copy numbers are not sufficiently reliable. Also, ASVs with low entropy (information contained in the time series) were excluded because reliable analyses of EDM require a sufficient amount of temporal information in the time series.

### Independent validations of the MiSeq sequencing with internal standard DNAs

The quantitative capacity of the MiSeq sequencing with internal standard DNAs (i.e., the quantitative MiSeq sequencing) is one of important factors that could influence subsequent data analyses. To check the reliability of the quantitative capacity of the method, I performed two independent DNA measurements (fluorescent-based total DNA quantifications and quantitative PCR [qPCR] of the 16S region) and compared the results with those of the quantitative MiSeq sequencing. Brief protocols of the experiments are described below. Details of the total DNA quantification, qPCR and shotgun metagenomic analysis and discussion of the results are provided in Supplementary Information.

Total DNAs were quantified using the Quant-iT assay kit (Promega, Madison, Wisconsin, USA). Three µl of each extracted DNA (from *ϕ* 0.22-*μ*m Sterivex) was mixed with the fluorescent reagent and DNA concentration was measured following the manufactuer’s protocol. The results were compared with the total DNA copy numbers estimated by quantitative MiSeq sequencing of four marker regions (i.e., 16S, 18S, ITS and COI) (Fig. S3C). An assumption behind the analysis is that most cellular organisms were captured by sequencing the four marker regions.

qPCR of the 16S region was performed using the same primer set used for the quantitative MiSeq sequencing (515F-806R primers) (Bates *et al*. 2011). Briefly, 2 *μ*l of each extracted DNA (from *ϕ* 0.22-*μ*m Sterivex) was added to an 8-*μ*l qPCR reaction containing 1 *μ*l of 5 *μ*M 515F primer, 1 *μ*l of 5 *μ*M 806R primer, 5 *μ*l of Platinum SuperFi PCR Master Mix (ThermoFisher Scientific, Waltham, Massachusetts, USA), 0.5 *μ*l of 20 × EvaGreen (Biotium, San Francisco, California, USA), and 0.5 *μ*l of MilliQ water. Sixty cycles of PCR were performed and the fluorescence was measured by LightCycler 480 (Roche, Basel, Switzerland). The total 16S copy numbers measured by qPCR correlated well with those measured by the quantitative MiSeq sequencing (Fig. S3D). In contrast, those measured by qPCR did not show a linear relationship with sequence reads (Fig. S3E).

Furthermore, to check whether and how PCR-based assessments of community composition biased the results, shotgun-metagenomic analysis was performed for a subset of the samples. Only four samples, of which the community diversity was high, were analyzed because much deeper sequencing is necessary for the shotgun metagenomic analysis. Briefly, approximately 10–30 ng of total DNA were used as inputs, and Illumina DNA Prep (Illumina, San Diego, CA, USA) was used to prepare the library for the shotgun metagenome. The library was prepared by following the manufacture’s protocol. The double-stranded DNA concentration of the library was then adjusted to 4 nM and the DNA was sequenced on MiSeq using a MiSeq V2 Reagent kit for 2 × 250 bp PE (Illumina, San Diego, CA, USA). In total, 20,601,323 reads (10 Gb for 4 samples) were generated. The low quality reads and adapter sequences were removed using fastp (Chen *et al*. 2018), and the filtered sequences were analyzed using phyloFlash (Gruber-Vodicka *et al*. 2020).

### Empirical dynamic modeling: Convergent cross mapping (CCM)

The reconstruction of the original dynamics using time-lagged coordinates is known as State Space Reconstruction (SSR) (Takens 1981; Deyle & Sugihara 2011) and is useful when one wants to understand complex dynamics. Recently developed tools for nonlinear time series analysis called “Empirical Dynamic Modeling (EDM),” which were specifically designed to analyze state-dependent behavior of dynamic systems, are rooted in SSR (Sugihara *et al*. 2012; Ye *et al*. 2015b; Deyle *et al*. 2016; Ushio *et al*. 2018a). These methods do not assume any set of equations governing the system, and thus are suitable for analyzing complex systems, for which it is often difficult to make reasonable *a priori* assumptions about their underlying mechanisms. Instead of assuming a set of specific equations, EDM recovers the dynamics directly from time series data, and is thus particularly useful for forecasting ecological time series, which are otherwise often difficult to forecast.

To detect causation between species detected by the DNA analysis, I used convergent cross mapping (CCM) (Sugihara *et al*. 2012; Osada & Ushio 2021). An important consequence of the SSR theorems is that if two variables are part of the same dynamical system, then the reconstructed state spaces of the two variables will topologically represent the same attractor (with a one-to-one mapping between reconstructed attractors). Therefore, it is possible to predict the current state of a variable using time lags of another variable. We can look for the signature of a causal variable in the time series of an effect variable by testing whether there is a correspondence between their reconstructed state spaces (i.e., cross mapping). This cross-map technique can be used to detect causation between variables. Cross-map skill can be evaluated by either a correlation coefficient (*ρ*), or mean absolute error (MAE) or root mean square error (RMSE) between observed values and predictions by cross mapping.

In the present study, cross mapping from one variable to another was performed using simplex projection (Sugihara & May 1990). How many time lags are taken in SSR (i.e, optimal embedding dimension; *E*) is determined by simplex projection using RMSE as an index of forecasting skill. More detailed algorithms about simplex projection and cross mapping can be found in previous reports (Sugihara & May 1990; Sugihara *et al*. 2012).

When the causal relationships between network properties were examined, I considered the interaction time lag between the network properties. This can be done by using “lagged CCM” (Ye *et al*. 2015a). For normal CCM, correspondence between reconstructed state space (i.e., cross-mapping) is checked using the same time point. In other words, information embedded in an effect time series at time *t* may be used to predict the state of a potential causal time series at time *t*. This idea can easily be extended to examine time-delayed influence between time series by asking the following question: is it possible to predict the state of a potential causal time series at time *t*–*tp* (*tp* is a time delay) by using information embedded in an effect time series at time *t*? Ye *et al*. (2015a) showed that lagged CCM is effective for determining the effective time delay between variables. In the present study, I examined the time delay of the effects from 0 to 14 days. When examining species interaction in the rice plots, the time delay of the interactions was fixed as –1 in order to avoid extremely large computational costs (i.e., 1197 × 1197 CCMs must be performed for each *tp*).

The significance of CCM is judged by comparing convergence in the cross-map skill of Fourier surrogates and original time series. More specifically, first, 1000 surrogate time series for one original time series are generated. Surrogate time series were generated so that they conserve seasonality (i.e., rEDM::make_surrogate_data(method = “seasonal”); details are in the scripts deposited). Five rice plot replicates were treated as if they were taken in five different years. Second, the convergence of the cross-map skill is calculated for these 1000 surrogate time series and the original time series. Specifically, the convergence of the cross-map skill (measured by ΔRMSE in the present study) is calculated as the cross-map skill at the maximum library length minus that at the minimum library length (Sugihara *et al*. 2012; Osada & Ushio 2021). Based on consideration of a large number of CCMs among 1197 DNA species, I used *P* = 0.005 as threshold. For CCMs among the network properties, I used *P* = 0.05, a more commonly used threshold.

### Empirical dynamic modeling: Multivariate, regularized S-map method

The multivariate S-map (sequential locally weighted global linear map) method allows quantifications of dynamic (i.e., time-varying) interactions (Sugihara 1994; Deyle *et al*. 2016). Consider a system that has *E* different interacting variables, and assume that the state space at time *t* is given by *x*(*t*) = {*x*_1_(*t*), *x*_2_(*t*), …, *x*_*E*_(*t*)}. For each target time point *t*^*^, the S-map method produces a local linear model that predicts the future value *x*_1_(*t*^*^ + *p*) from the multivariate reconstructed state space vector *x*(*t*^*^). That is,

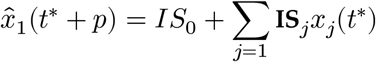

where 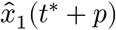 is a predicted value of *x*_1_ at time *t*^*^ + *p*, and *IS*_0_ is an intercept of the linear model. The linear model is fit to the other vectors in the state space. However, points that are close to the target point, *x*(*t*^*^), are given greater weighting (i.e., locally weighted linear regression). Note that the model is calculated separately for each time point, *t*. As recently shown, **IS**_*j*_, the coefficients of the local linear model, are a proxy for the interaction strength between variables (Deyle *et al*. 2016).

In the present study, all ASVs that have causal influences on a focal species (determined by CCM) and have non-zero abundance at a target time point were included in the multivariate, regularized S-map. In some cases, the number of causal species exceed the optimal *E*. In that case, I simply added all the detected causal species in the S-map. Nonetheless, in the present study, the number of variables (= the number of causal species for a target species at each time point) included in each S-map model was fewer than 14 for most target species (over 99% of all cases; see Fig. S5B), which allowed rigorous estimations of interaction strengths between species (i.e., the number of data point, 610, is greater than the square of the embedding dimension). In the same way as with simplex projection and CCM, the performance of the multivariate S-map was also measured by RMSE (or a correlation coefficient, *ρ*) between observed and predicted values by the S-map (i.e., leave-one-out cross validation). In the present study, to reduce the possibility of overestimation and to improve forecasting skill, a regularized version of multivariate S-map was used (Cenci *et al*. 2019).

### Calculations of properties of the interaction network

Properties of the reconstructed interaction network calculated include: ASV diversity, the number of interactions, connectance, mean interaction strength (*IS*) per link, mean interaction capacity, dynamic stability and coefficient of variation (CV) in population dynamics. Next, I give the definitions of the properties.

ASV diversity and the number of interactions are the number of ASVs present in a community and the number of interactions among ASVs present in a community, respectively. The existence of interactions was defined by significant CCM results. In addition, even if CCM detected significant interactions between two species, the interaction at a certain time point was judged absent if either or both of the species was/were absent at the time point. Connectance, *C*, is defined as *C* = *N*_*link*_ / *S*^2^, where *S* and *N*_*link*_ indicate the number of species and the number of interactions (links) in a community, respectively. Mean interaction strength per link, *IS*_*link*_, was calculated as follows:

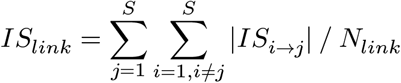

where *IS*_*i*→*j*_ indicates an S-map coefficient from *i*th species to *j*th species. Note that I took the absolute value of the S-map coefficient when calculating *IS*_*link*_. Mean interaction capacity, *IC*, was calculated as follows:

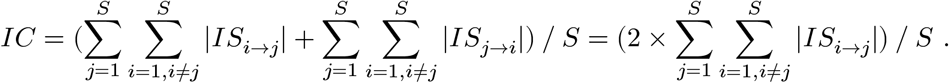

Species interaction capacity is defined as the sum of interactions that a single species gives and receives, and mean interaction capacity of a community is the averaged species interaction capacity. Dynamic stability of the community dynamics was calculated as the absolute value of the dominant eigenvalue of the interaction matrix (i.e., local Lyapunov exponent) as described in a previous study (Ushio *et al*. 2018b). CV of the community dynamics at time *t* was calculated as follows:

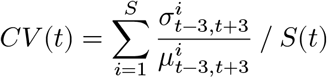

where 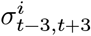 and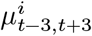 indicate the standard deviation and mean value of the abundance of species *i* from time *t*–3 to *t* + 3, respectively (i.e., one-week time window). *S*(*t*) is the number of ASVs at time *t*.

### Empirical dynamic modeling: Random shuffle surrogate test

To test whether the patterns generated (e.g., in Fig. 3) are statistical artifacts, I did a random shuffle surrogate test. In the test, the original time series were randomly shuffled within a plot using the rEDM::make_surrogate_shuffle() function in the rEDM package (Ye *et al*. 2015b, 2018) of R. Then, the same number of causal pairs was randomly assigned in a randomly shuffled ecological community. The regularized, multivariate S-map and subsequent analyses of the network properties (all identical to the original analyses) were applied to the randomly shuffled time series.

### Meta-analysis of biodiversity, temperature and abundance

To validate my hypothesis that the diversity is determined by interaction capacity and connectance, and that they are influenced by temperature and total organism abundance, I compiled published data from various ecosystems. The collected data include two global datasets and four local datasets collected in Japan: (1) global ocean microbes (Sunagawa *et al*. 2015), (2) global soil microbes (Bahram *et al*. 2018), (3) fish from a coastal ecosystem (Masuda 2008), (4) prokaryotes from freshwater lake ecosystems (Okazaki *et al*. 2017), (5) zooplankton from a freshwater lake ecosystem (Sakamoto *et al*. 2018) and (6) benthic macroinvertebrates from freshwater tributary lagoon ecosystems (Okano *et al*. 2018). Because the influences of temperature and total species abundance/biomass on community diversity (or interaction capacity and connectance) are likely to be nonlinear, I adopted a general additive model (Wood 2004) as follows:

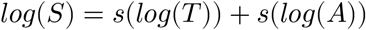

where *S, T, A* and *s*() indicate species diversity (or OTU diversity), temperature, an index of total species abundance (or biomass) and a smoothing term, respectively. The relationships between diversity, temperature and total abundance were analyzed using the model described in the main text. GAM was performed using the “mgcv” package of R (Wood 2004).

Data analyzed in the meta-analysis were collected from the publications or official websites (Sunagawa *et al*. 2015; Sakamoto *et al*. 2018), or provided by the authors of the original publications (Masuda 2008; Okazaki *et al*. 2017; Bahram *et al*. 2018; Okano *et al*. 2018). Therefore, raw data for the meta-analysis are available from the original publications, or upon reasonable requests to corresponding authors of the original publications. Scripts for the meta-analysis are available in Github (https://github.com/ong8181/interaction-capacity).

### Computation and results visualization

Simplex projection, S-map and CCM were performed using “rEDM” package (version 0.7.5) (Ye *et al*. 2015b, 2018), with results visualized using “ggplot2” (Wickham 2009). Data were analyzed in the free statistical environment R3.6.1 (R Core Team 2019).

## Data and code availability

All scripts used in the present study are available in Github (https://github.com/ong8181/interaction-capacity). Sequence data are deposited in DDBJ Sequence Read Archives (DRA) (DRA accession number = DRA009658, DRA009659, DRA009660 and DRA009661 for ecological community monitoring data and DRA012505 for supporting shotgun metagenomic data), which will be publicly available after the formal acceptance of the preprint.

## Acknowledgements

I thank Asako Kawai for the assistance in field monitoring and DNA library preparations, Saori Furukawa and Atsushi J. Nagano for help in the additional DNA experiments, Akira Matsumoto, Satoru Yonezawa, and Satomi Yoshinami for assistance in the monitoring setup and field monitoring, Yutaka Osada for advice and discussion on empirical dynamic modeling, Takeshi Miki, Erik A. Hobbie, Masahiro Ryo, and Hao Ye for comments on the manuscript, and Ai Matsuda for help in the figure editing. I thank Mohammad Bahram, Yukiko Goda, Jun-ichi Okano, Yusuke Okazaki, Noboru Okuda, and Jun-ya Shibata for providing the data set for the meta-analysis. I also thank Christoph Ratzke and Akira Yamawo for providing the raw data published in their previous studies. This research was supported by PRESTO (JPMJPR16O2) from the Japan Science and Technology Agency (JST) and the Hakubi Project in Kyoto University.

## Author Contributions

M.U. conceived the idea, designed research, collected data, analyzed DNA sequences and ecological time series, performed meta-analysis and wrote the manuscript.

