## Supplementary Figures for "Interaction capacity underpins community diversity"

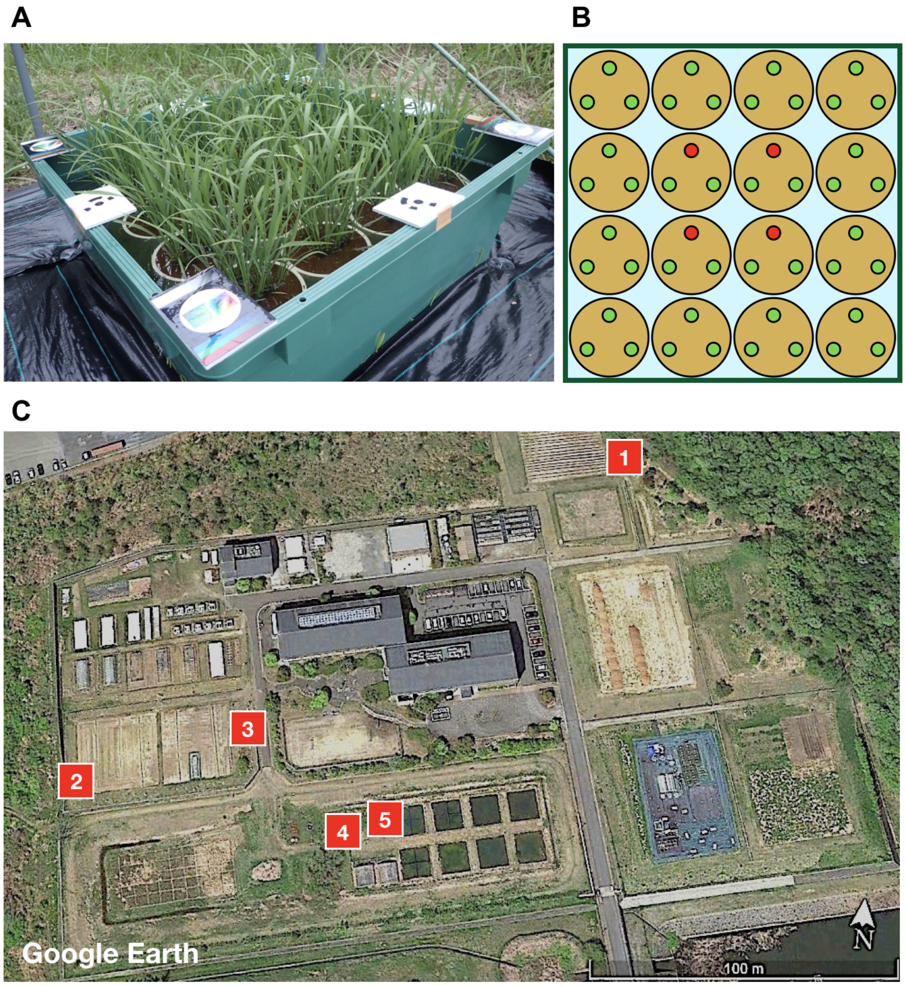


**Figure S1. Experimental rice plots established in Center for Ecological Research, Kyoto Univeristy (34° 58'18' ' N, 135° 57'33' ' E) in 2017.** **A** An image of the experimental rice plot (90 cm × 90 cm; the photo was taken by M. Ushio). **B** Three rice individuals were grown in each Wagner pot. Growth rates of four individuals at the center of each plot were monitored for agricultural purposes. The growth data are being analyzed for different purposes and thus are not shown in this study. **C** Five experimental rice plots were established in an experimental field in Center for Ecological Research, Kyoto University (the image taken from Google Earth). Red squares indicate the locations of the rice plots.

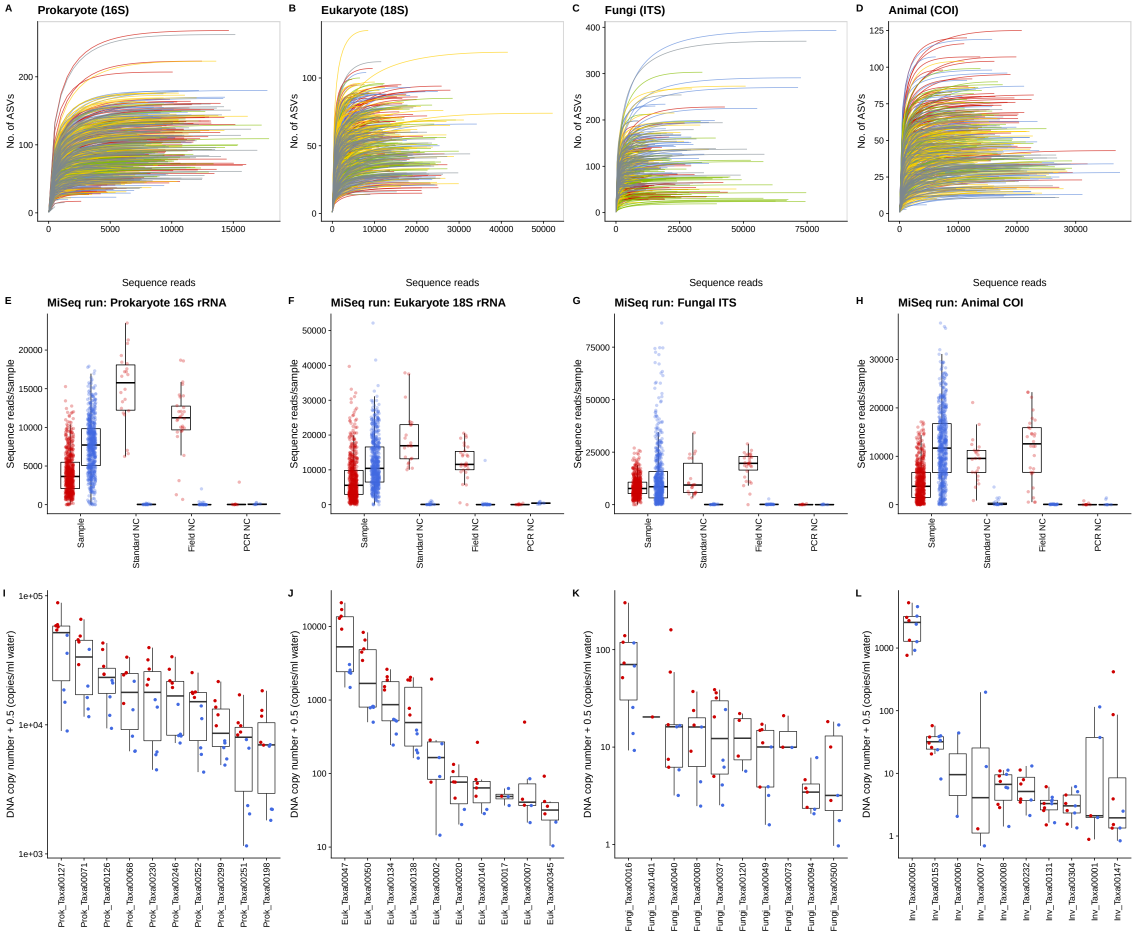


**Figure S2. Rarefaction curve and results of negative and positive controls.** Rarefaction curve (**A**-**D**) and the results of analysis of negative (**E**-**H**) and positive (**I**-**L**) controls. Most of the rarefaction curves reached a plateau, suggesting that the sequencing captured most diversity of the target taxa (**A**-**D**). Investigations of negative control samples suggested that no serious contamination occurred during sample transportation, eDNA extraction or library preparation (Almost no non-standard DNAs were detected from Standard NC, Field NC or PCR NC; **E**-**H**). No dramatic reductions in DNA copy numbers of the top 10 dominant taxa were observed in samples from which DNAs were extracted before the storage duration exceeded 3 months (red points in **I**-**L**; note that DNAs of all samples analyzed in the time series analysis were extracted within 1-2 months after sampling). The box-plot elements are defined as follows: center line, median; box limits, upper and lower quantiles; whiskers, 1.5 $\times$ interquantile range; points, outliers. Together, these results suggest that the water sampling and DNA analysis in the present study enabled reliable illustrations of the dynamics of ecological communities in the rice plots. Detailed descriptions of the negative and positive control samples are provided in the Supplementary Text.

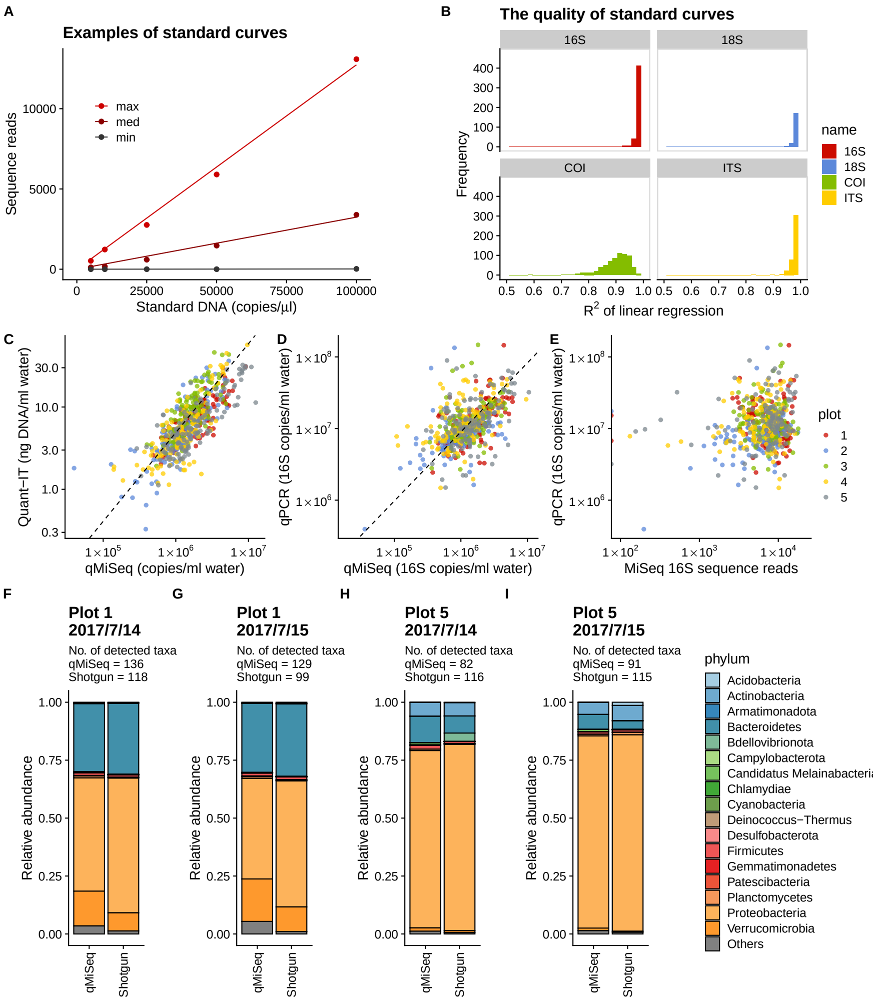


**Figure S3. Validation of the quantitative MiSeq sequenicing ("qMiSeq"). A** Examples of the relationship between standard DNA copy numbers and sequence reads. Slope of the linear regression changes among samples, partly because of the different intensity of PCR inhibition. The regression lines with maximum, medium, and minimum slope values are shown by red, dark red, and black points and lines as examples. **B** Distributions of $R^2$ values of the linear regression (610 samples $\times$ 4 marker regions). Most samples and regions show $R^2 > 0.9$, suggesting that the sequence reads for each standard DNA are proportional to the initial DNA copy numbers. **C** The relationship between total DNA concentrations quantified by fluorescent-based method and the total DNA copy numbers estimated by qMiSeq. **D** The relationship between total 16S copy numbers measured by quantitative PCR (qPCR) and qMiSeq. **E** The relationship between total 16S copy numbers measured by qPCR and 16S sequence reads. Dashed line and colors indicate a linear regression and the rice plots, respectively. **F-I** Comparisons of 16S amplicon sequencing (qMiSeq) and 16S extracted from the results of shotgun metagenomic analysis by the phyloFlash pipeline. Barplots indicate community composition revealed by the two methods. The number of taxa detected by each method is shown at the top of the barplots. Because the shotgun metagenomic analysis requires much deeper sequencing than an amplicon sequencing, only four samples were analyzed for the comparison. Samples were from **F** Plot 1 on 2017/7/14, **G** Plot 1 on 2017/7/15, **H** Plot 5 on 2017/7/14, and **I** Plot 5 on 2017/7/15. Detailed methods and discussions are presented in Supplementary Information.

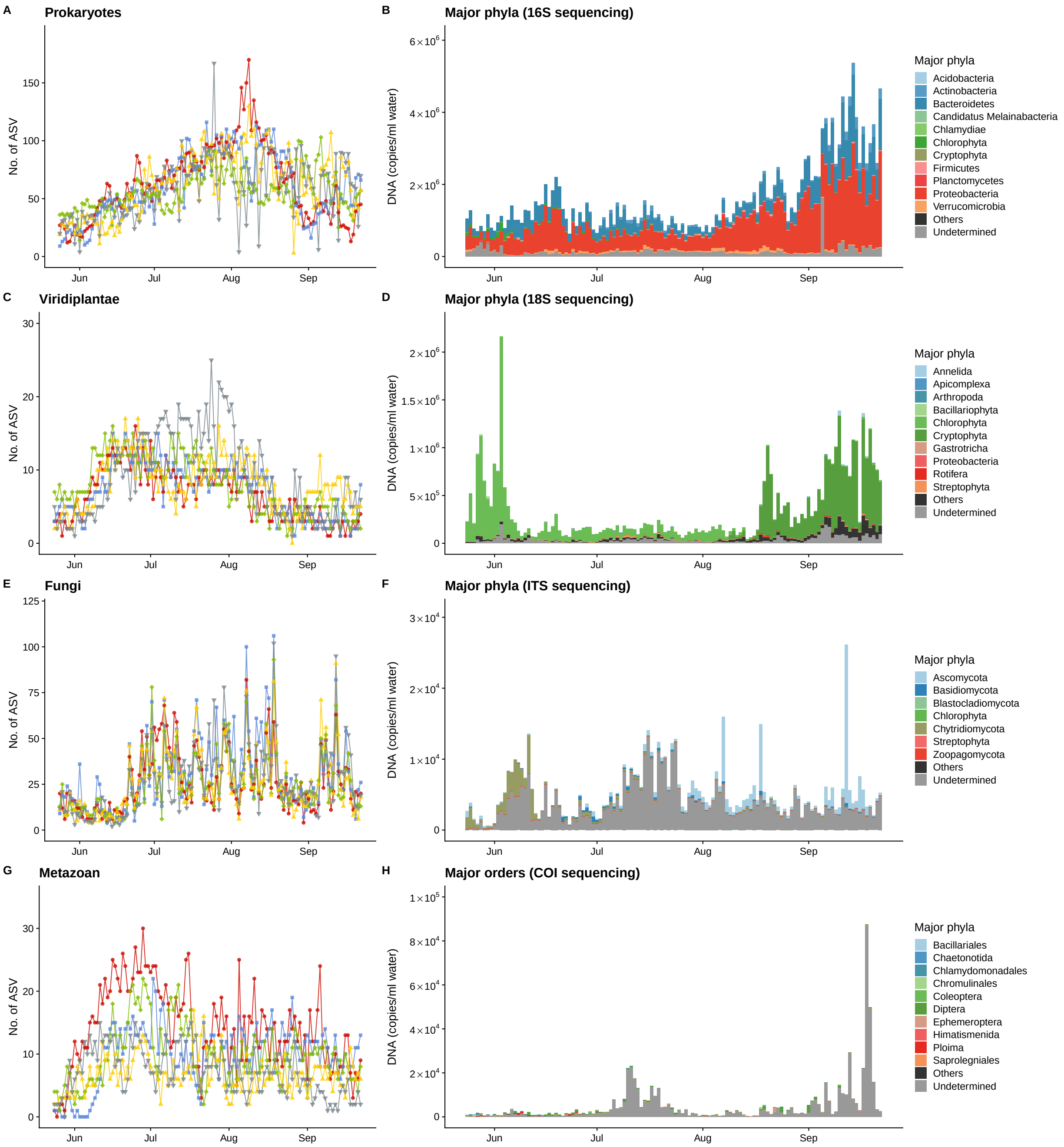


**Figure S4. Patterns in DNA time series of major taxonomic groups in the study.** Temporal patterns of ASV diversity, major taxa and the DNA copy numbers for **A-B** prokaryotes, **C-D** fungi, **E-F** metazoans, and **G-H** viridiplantae.

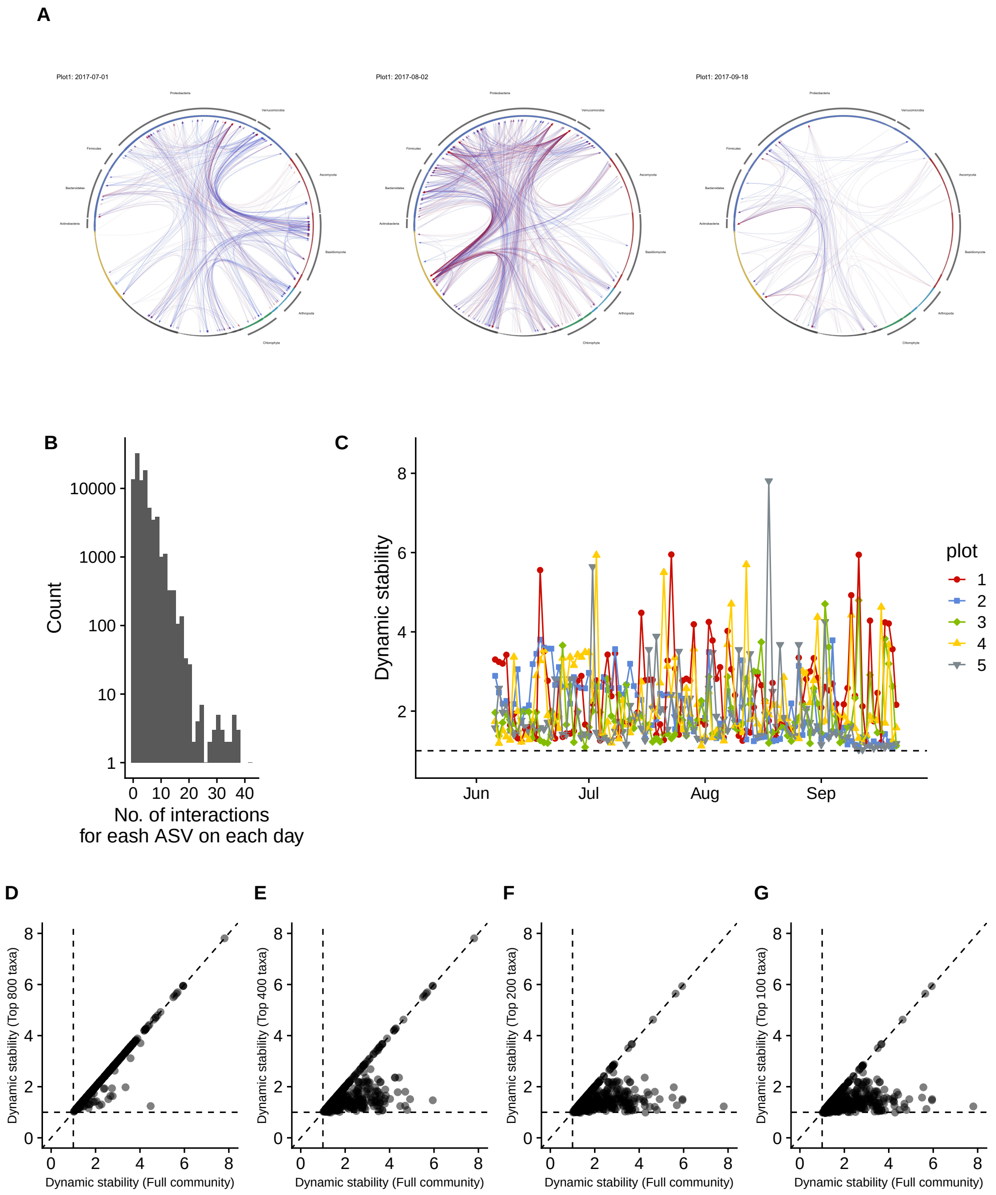


**Figure S5. Time-varying properties of the interaction network and dependence of dynamic stability on the community size. A** Changes in the network structures. Interaction networks in Plot 1 on 1 July, 2 August and 18 September 2017 are shown as examples of time-varying interaction networks. Blue and red arrows indicate positive and negative causal interactions, respectively. **B** The number of interactions that each ASV had on each day. Note that $y$-axis is log-transformed. **C** Dynamic stability of the ecological community. Different colors, lines and symbols indicate different plots. **D**-**G** Dependence of dynamic stability on the community size. The scatter plots comparing the dynamic stability of the full community (1197 ASVs) and the most dominant 800 (**D**), 400 (**E**), 200 (**F**), and 100 (**G**) ASVs are shown. Dashed diagonal line indicates 1:1 line. Dashed horizontal and vertical lines indicate the dynamic stability value 1 (values lower than 1 indicate locally stable dynamics). As shown in **C**-**F**, the removals of moderately abundant ASVs stabilize the community dynamics, suggesting that fluctuations in the abundance of the moderately abundant ASVs could contribute to the unstable community dynamics in the rice plots.

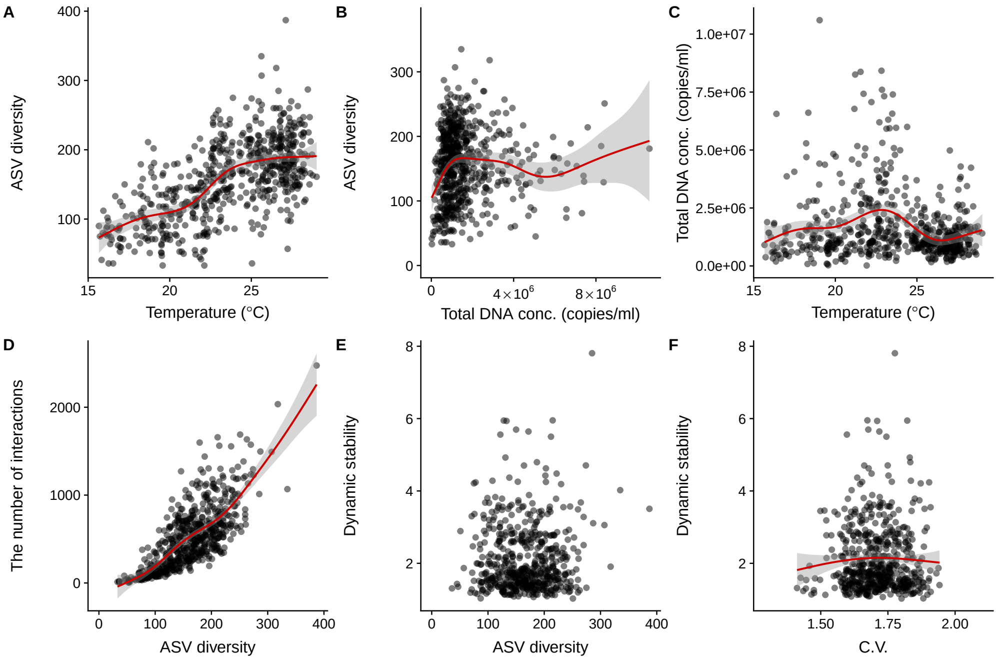


**Figure S6. Additional patterns in network properties.** **A** Mean air temperature and ASV diversity, **B** total DNA copy numbers and ASV diversity, **C** Mean air temperature and total DNA copy numbers, **D** The number of interactions and ASV diversity. **E** ASV diversity and dynamic stability and **F** Dynamic stability and coefficients of variation of community dynamics. Red lines indicate statistically clear nonlinear regressions by the general additive model (GAM). Gray shaded region indicates 95% confidence interval of GAM.

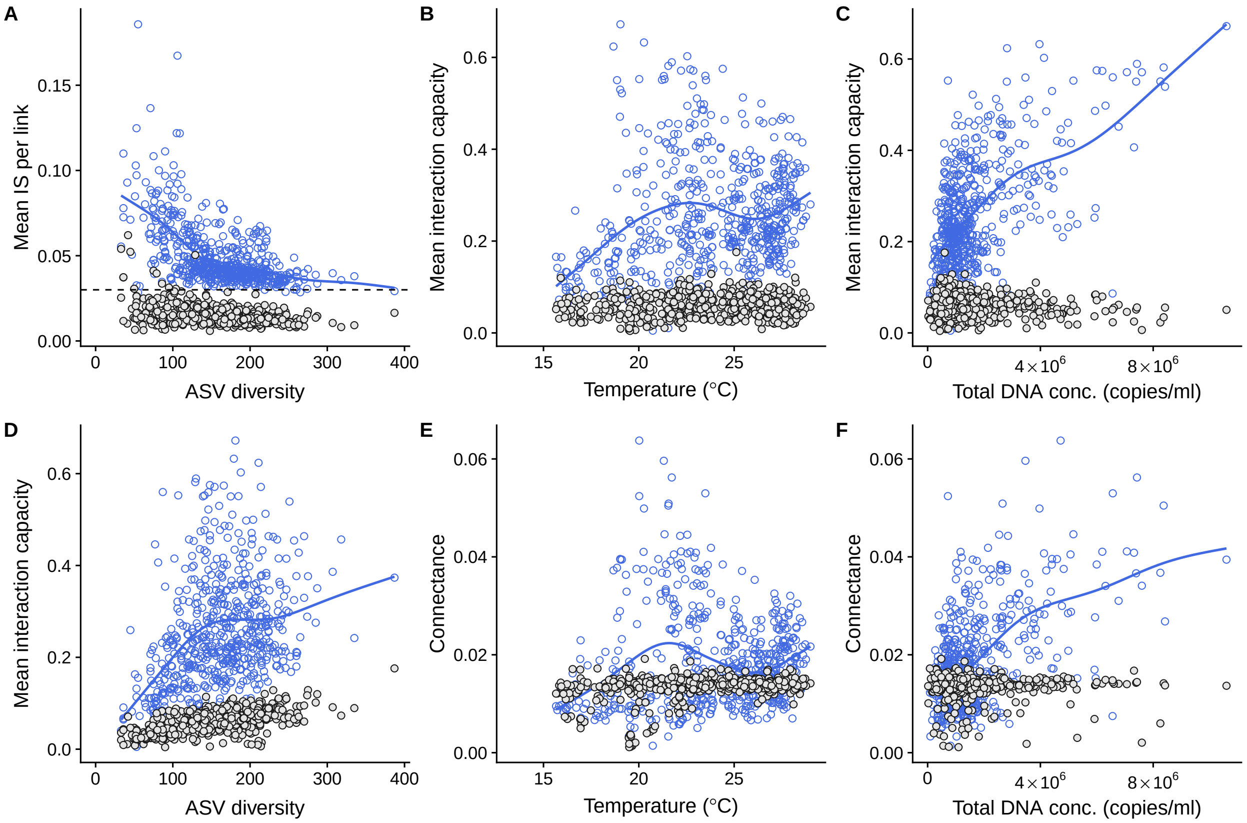


**Figure S7. Comparisons between observed network properties and those of randomly shuffled time series.** **A** Mean interaction strength and ASV diversity, **B**, **C** Mean interaction capacity, mean air temperature and total DNA copy numbers, and **D** mean interaction capacity and ASV diversity, and **E**, **F** connectance, mean air temperature and total DNA copy numbers. Gray points indicate patterns generated by analyzing randomly shuffled time series. Blue points and line indicate patterns for the original time series. The observed patterns were not reproduced by the analyses of the randomly shuffled time series.

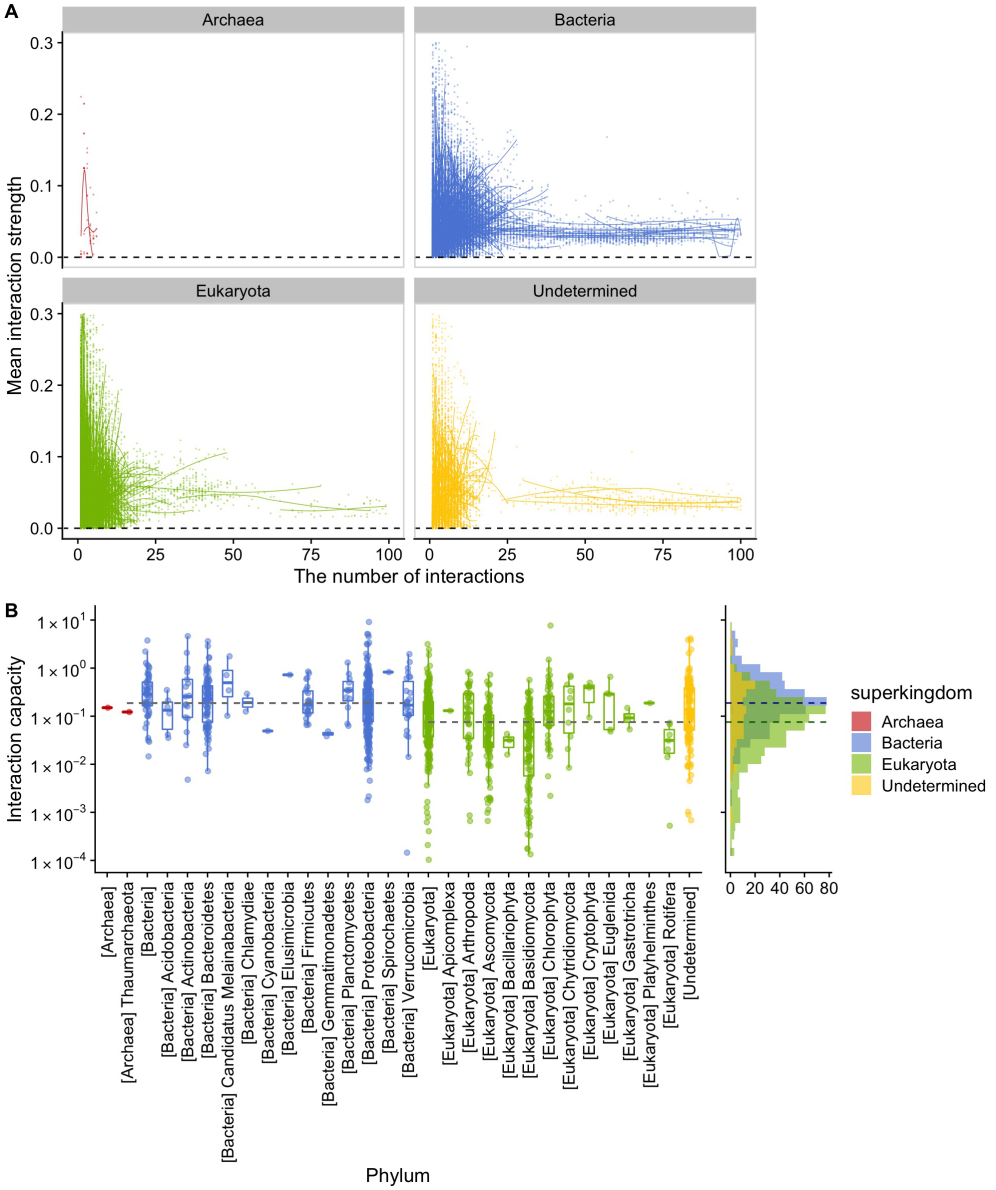


**Figure S8. Species-level patterns of interaction strength and capacity.** **A** Relationships between the number of interactions and mean interaction strength per link. **B** Relationship between species-level interaction capacity and phylum. Species-level interaction capacity can be calculated for each day in each plot, and the median value of the interaction capacity is plotted as a representative interaction capacity for each ASV in **B**. The box-plot elements are defined as follows: center line, median; box limits, upper and lower quantiles; whiskers, 1.5 × interquantile range; points, outliers. Dashed lines indicate the median interaction capacity for Bacteria and Eukaryota. Bacterial taxa generally show a higher interaction capacity than eukaryotes, suggesting that the potential maximum diversity of Bacteria is higher than that of Eukaryota.

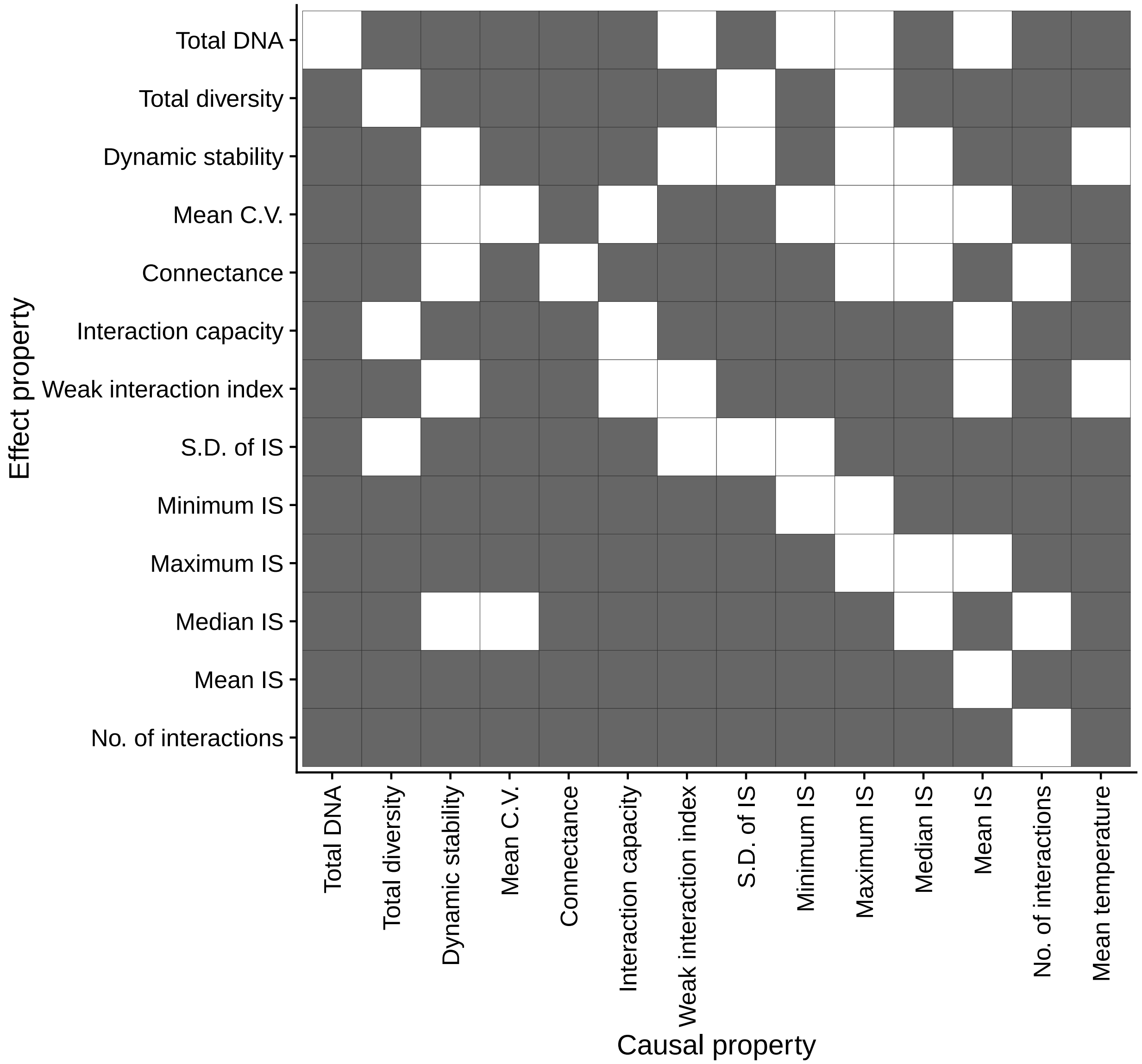


**Figure S9. Causal relationships among the network properties.** A cell filled with gray color indicates that there is a causal influence from a column variable ("Causal property") to a row variable ("Effect property"), which is detected by convergent cross mapping.

### Table S1. Statistical results of GAM between network properties.

| Explained variable | Smooth term (explaining variable) | Effective d.f. | *F* value | *P* value | Adjusted $R^2$ |
|---|---|---|---|---|---|
| Connectance | Total ASV diversity | **4.7** | **6.4** | **$2.35 \times 10^{-6}$** | **0.06** |
| Connectance | Mean air temperature | **4.8** | **8.4** | **$1.55 \times 10^{-8}$** | **0.079** |
| Connectance | Total DNA concentrations | **5.1** | **42.9** | **$2.68 \times 10^{-48}$** | **0.319** |
| Dynamic stability | Mean C.V. | 1.8 | 0.8 | 0.426 | 0.002 |
| Dynamic stability | Total ASV diversity | **6.5** | **2.7** | **$9.91 \times 10^{-3}$** | **0.029** |
| Interaction capacity | Total ASV diversity | **3.7** | **24.6** | **$3.21 \times 10^{-21}$** | **0.169** |
| Interaction capacity | Mean air temperature | **4.3** | **10.8** | **$3.36 \times 10^{-10}$** | **0.089** |
| Interaction capacity | Total DNA concentrations | **6.9** | **40.2** | **$6.01 \times 10^{-57}$** | **0.356** |
| Interaction capacity | Connectance | **6.1** | **177.9** | **$1.79 \times 10^{-235}$** | **0.694** |
| Maximum IS | Total ASV diversity | 3.2 | 1 | 0.404 | 0.005 |
| Maximum IS | Mean air temperature | 1 | 2.7 | 0.0983 | 0.003 |
| Mean C.V. | Total ASV diversity | **4.8** | **10.8** | **$2.34 \times 10^{-11}$** | **0.094** |
| Mean IS | Total ASV diversity | **5.4** | **52.5** | **$1.30 \times 10^{-61}$** | **0.376** |
| No. of interactions | Total ASV diversity | **5** | **129.5** | **$9.13 \times 10^{-146}$** | **0.581** |
| Total ASV diversity | Total DNA concentrations | **6.5** | **4.9** | **$1.29 \times 10^{-5}$** | **0.054** |
| Total DNA concentrations | Mean air temperature | **5.8** | **10.6** | **$1.42 \times 10^{-12}$** | **0.107** |
