## Supplementary Text for "Interaction capacity underpins community diversity"

### Supplementary Information for "Interaction capacity underpins community diversity"

Masayuki Ushio

#### Contents:

1. Thermal cycle profiles
2. Internal standard DNA sequences
3. Summary of sequence quality
4. Rarefaction curve
5. Negative control results
6. Positive control results
7. Total DNA quantification
8. Quantitative PCR (qPCR)
9. Shotgun metagenomic analysis
10. Software packages
11. Sample meta-data (separete xlsx file)
12. Supplementary video

### 1. Thermal cycle profiles

#### Common protocols

The first-round PCR (first PCR) was carried out with a 12-$\mu$l reaction volume containing 6.0 $\mu$l of 2 $\times$ KAPA HiFi HotStart ReadyMix (KAPA Biosystems, Wilmington, WA, USA), 0.7 $\mu$l of forward primer, 0.7 $\mu$l of reverse primer (each primer at 5 $\mu$M used in the reaction; with adaptor and six random bases), 2.6 $\mu$l of MilliQ water, 1.0 $\mu$l of DNA template and 1.0 $\mu$l of internal standard DNA mixture. Thermal cycle profile, primer set and internal standard DNA sequences are provided in following subsections. Triplicate first PCRs were performed, and these replicate products were pooled in order to mitigate the PCR dropouts. The pooled first PCR products were purified using AMPure XP (PCR product : AMPure XP beads = 1:0.8; Beckman Coulter, Brea, California, USA). The pooled, purified, and 10-fold diluted first PCR products were used as templates for the second-round PCR.

The second-round PCR (second PCR) was carried out with a 24-$\mu$l reaction volume containing 12 $\mu$l of 2 $\times$ KAPA HiFi HotStart ReadyMix, 1.4 $\mu$l of each primer (each primer at 5 $\mu$M in the reaction volume), 7.2 $\mu$l of MilliQ water and 2.0 $\mu$l of template. Different combinations of forward and reverse indices were used for different templates (samples) for massively parallel sequencing with MiSeq. The thermal cycle profile after an initial 3 min denaturation at 95°C was as follows (12 cycles): denaturation at 98°C for 20 s; annealing at 68°C for 15 s; and extension at 72°C for 15 s, with a final extension at 72°C for 5 min.

Twenty microliters of the indexed second PCR products were mixed, and the combined library was again purified using AMPure XP (PCR product: AMPure XP beads = 1:0.8). Target-sized DNA of the purified library (*ca.* 440 bp for prokaryote 16S rRNA; *ca.* 320 bp for eukaryote 18S rRNA; *ca.* 470-600 bp for fungal ITS; *ca.* 510 bp for animal COI) was excised using E-Gel SizeSelect (ThermoFisher Scientific, Waltham, MA, USA) (Note that "the target size" includes Illumina P5/P7 adapter, Rd1SP/Rd2SP sequencing primer and sample-specific index sequences of which total length is 137 bp). The double-stranded DNA concentration of the library was quantified using a Qubit dsDNA HS assay kit and a Qubit fluorometer (ThermoFisher Scientific, Waltham, MA, USA). The double-stranded DNA concentration of the library was then adjusted using MilliQ water and the DNA was applied to the MiSeq (Illumina, San Diego, CA, USA). The prokaryote 16S rRNA, eukaryote 18S rRNA, fungal ITS and animal COI libraries were sequenced using MiSeq Reagent Kit V2 for 2 $\times$ 250 bp PE, MiSeq Reagent Kit V2 for 2 $\times$ 150 bp PE, MiSeq Reagent Kit V3 for 2 $\times$ 300 bp PE and MiSeq Reagent Kit V2 for 2 $\times$ 250 bp PE, respectively.

#### Primer sequences

For the first PCR, taxa-specific univarsal primers were combined with the MiSeq sequencing primers and six random bases (Ns) to improve the quality of MiSeq sequencing. For the second PCR, MiSeq adaptor and sequencing primers were combined with index sequences (eight bases denoted by X in the following table) to identify each sample. See sample metadata for index sequences of each sample.

**Table S2. Taxa-specific 1st PCR primers**

| Target taxa | Primer name | Sequencing primer - NNNNNN - Universal primer |
|---|---|---|
| Prokaryote (16S rRNA) | 515F | TCGTCGGCAGCGTCAGATGTGTATAAGAGACAG NNNNNN GTGYCAGCMGCCGCGGTAA |
| | 806R | GTCTCGTGGGCTCGGAGATGTGTATAAGAGACAG NNNNNN GGACTACNVGGGTWTCTAAT |
| Eukaryote (18S rRNA) | Euk_1391f | TCGTCGGCAGCGTCAGATGTGTATAAGAGACAG NNNNNN GTACACACCGCCCGTC |
| | EukBr | GTCTCGTGGGCTCGGAGATGTGTATAAGAGACAG NNNNNN TGATCCTTCTGCAGGTTCACCTAC |
| Fungi (ITS) | ITS1_F_KYO1 | TCGTCGGCAGCGTCAGATGTGTATAAGAGACAG NNNNNN CTHGGTCATTTAGAGGAASTAA |
| | ITS_KYO2 | GTCTCGTGGGCTCGGAGATGTGTATAAGAGACAG NNNNNN TTYRCTRCGTTCTTCATC |
| Animal (mitocondoria COI) | mlCOIintF | ACACTCTTTCCCTACACGACGCTCTTCCGATCT NNNNNN GGWACWGGWTGAACWGTWTAYCCYCC |
| | HCO2198 | GTGACTGGAGTTCAGACGTGTGCTCTTCCGATCT NNNNNN TAAACTTCAGGGTGACCAAAAAATCA |

References for the taxa-specific 1st PCR primers are as follows: 515F-806R (Bates *et al.* 2011; Caporaso *et al.* 2011), Euk_1391f-EukBr (Amaral-Zettler *et al.* 2009), ITS1_F_KYO1-ITS_KYO2 (Toju *et al.* 2012), mlCOIintF (Leray *et al.* 2013) and HCO2198 (Folmer *et al.* 1994).

**Table S3. 2nd PCR primers**

| Primer name | | MiSeq adaptor - XXXXXXXX - Sequencing primer |
|---|---|---|
| 2nd PCR primer for 16S, 18S and ITS | Forward | AATGATACGGCGACCACCGAGATCTACAC XXXXXXXX TCGTCGGCAGCGTCAGATGTGTATAAGAGACAG |
| | Reverse | CAAGCAGAAGACGGCATACGAGAT XXXXXXXX GTCTCGTGGGCTCGGAGATGTGTATAAGAGACAG |
| 2nd PCR primer for COI | Forward | AATGATACGGCGACCACCGAGATCTACAC XXXXXXXX ACACTCTTTCCCTACACGACGCTCTTCCGATCT |
| | Reverse | CAAGCAGAAGACGGCATACGAGAT XXXXXXXX GTGACTGGAGTTCAGACGTGTGCTCTTCCGATCT |

**Taxa-specific thermal cycle profiles of the first-round PCR**

**Prokaryote 16S rRNA (515F-806R)**

The thermal cycle profile after an initial 3 min denaturation at 95°C was as follows (35 cycles): denaturation at 98°C for 20 s; annealing at 60°C for 15 s; and extension at 72°C for 30 s, with a final extension at the same temperature for 5 min.

**Eukaryote 18S rRNA (Euk_1391f-EukBr)**

The thermal cycle profile after an initial 3 min denaturation at 95°C was as follows (35 cycles): denaturation at 98°C for 20 s; annealing at 62°C for 15 s; and extension at 72°C for 30 s, with a final extension at the same temperature for 5 min.

**Fungal ITS (ITS1_F_KYO1-ITS_KYO2)**

The thermal cycle profile after an initial 3 min denaturation at 95°C was as follows (35 cycles): denaturation at 98°C for 20 s; annealing at 55°C for 15 s; and extension at 72°C for 30 s, with a final extension at the same temperature for 5 min.

**Animal COI (mlCOIintF-HCO2198)**

The thermal cycle profile after an initial 3 min denaturation at 95°C was as follows (35 cycles): denaturation at 98°C for 20 s; annealing at 55°C for 15 s; and extension at 72°C for 30 s, with a final extension at the same temperature for 5 min.

#### 2. Internal standard DNA sequences

Five artificially designed and synthesized internal standard DNAs, which are similar but not identical to the corresponding region of any existing target organisms (e.g., the V4 region of prokaryotic 16S rRNA), were included in the library preparation process to estimate the number of DNA copies [i.e., quantitative MiSeq sequencing; Ushio (2019); Ushio *et al.* (2018)]. They were designed to have the same primer-binding regions as those of known existing sequences and conserved regions in the insert region. Variable regions in the insert region were replaced with random bases so that no known existing sequences had the same sequences as the standard sequences. The numbers of standard DNA copies were adjusted appropriately to obtain a linear regression line between the copy numbers of the standard DNAs and their sequence reads from each sample. Sequences and the copy numbers of the standard DNAs are listed in the following tables.

Ushio et al. (2018) showed that the use of standard DNAs in MiSeq sequencing provides reasonable estimates of the DNA quantity in environmental samples when analyzing fish environmental DNA. Indeed, the copy numbers of standard DNAs were highly correlated with the number of sequence reads in this study (for most samples, $R^2 > 0.9$; Fig. S5B, suggesting that the number of sequence reads is generally proportional with the number of DNA copies in a single sample. However, it should be noted that it corrects neither for sequence-specific amplification efficiency, nor for species-specific DNA extraction efficiency. In other words, the method assumes similar amplification efficiencies across sequences, and similar DNA extraction efficiencies across microbial species, which is apparently not valid for complex environmental samples. Therefore, we need careful interpretations of results obtained using that method.

**Table S4. Sequences and the copy numbers of standard DNAs of prokaryote 16S rRNA**

| Primer name | Sequence |
|---|---|
| STD_pro1 | GTGCCAGCAGCCGCGGTAATACGTAGGGGGCAAGCGTTATCCGGAATTACTGGGCGTAAA |
| 5,000 copies/$\mu$l | GGGAAGGTAGGCGGAAGCTGAAGTCATGTGTGAAAACGCCTGGCTCAACTTAGCTCAGGG |
| | TCGCTGAAATCGTTGGATTCTCTGTGAACAAGGTCCTCGGAAATTTCTTGCTAGAGCAAT |
| | AAGGGCTGGCGGATAAGGCGAAACTTTGCGGCGAAGGCACGGAGCATAGGCAGGGCCCTA |
| | ACTGGACCGTTAAAGCGTGGGGAGCAAACAGGATTAGATACCCTGGTAGTCC |
| STD_pro2 | GTGCCAGCAGCCGCGGTAATACGTAGGGGGCAAGCGTTATCCGGAATTACTGGGCGTAAA |
| 10,000 copies/$\mu$l | GGGTATGTAGGCGGTAAAGGAAGTTACGAGTGAAATTACAGGGCTCAACCGATAAGTCGT |
| | GGCCAAGAGGCGGAGCCTGCAGTAATCGAGAGGTGGGCGGAATCGGGTAAAAGTCAACCT |
| | AAGGAAACAACTTTCAAAGGAAGCCAGATCGCGAAGGCGGCCTCGTAGTTAGGTCCTTCT |
| | CTGGACAAACGAAAGCGTGGGGAGCAAACAGGATTAGATACCCTGGTAGTCC |
| STD_pro3 | GTGCCAGCAGCCGCGGTAATACGTAGGGGGCAAGCGTTATCCGGAATTACTGGGCGTAAA |
| 25,000 copies/$\mu$l | GGGTTAGTAGGCGGGCAATTAAGTTATAGGTGAAAAGTAATGGCTCAACTTCGCGAACTC |
| | CGGACGCAGAAAGGTGCTGTGGCGTTAAAGGGGCTCCCGGAACCGAAGCAACAAACCTGA |
| | AAGCCGTTCCGATGTTTCGGAACTGCTAATGCGAAGGCTCAAGTCCTAGTAACTGGCGAT |
| | GTCCTAATCACAAAGCGTGGGGAGCAAACAGGATTAGATACCCTGGTAGTCC |
| STD_pro4 | GTGCCAGCAGCCGCGGTAATACGTAGGGGGCAAGCGTTATCCGGAATTACTGGGCGTAAA |
| 50,000 copies/$\mu$l | GGGAGTGTAGGCGGCCACGTAAGTTCCCGGTGAAATCGAGCGGCTCAACGTGCTCCTCGC |
| | CCGAGGTATCTAAGCCTTGTAACTCAGCCTAGGAACCCGGAAAGACTTTGTAGACATGCT |
| | AAAACGTGACCTGGTGTATGAATATCTCCAGCGAAGGCCGCACAACCGCACTCCACGCTT |
| | TAGTGGGTTTAAAAGCGTGGGGAGCAAACAGGATTAGATACCCTGGTAGTCC |
| STD_pro5 | GTGCCAGCAGCCGCGGTAATACGTAGGGGGCAAGCGTTATCCGGAATTACTGGGCGTAAA |
| 100,000 copies/$\mu$l | GGGCCGGTAGGCGGCTCCCGAAGTCGGTAGTGAAATTCTGAGGCTCAACTAAAACCACGA |
| | CATTGGAACGAGGTGTGTTAGCTTGATACCCGGCCAGTGGAACAGTGCTGCTCACTTCAT |
| | AACTCGGGTACGAAATAAGGAAGACGTCCCGCGAAGGCTTTGGCAGGGCATGCCAGGGCA |
| | TGGCTTTTGTTAAAGCGTGGGGAGCAAACAGGATTAGATACCCTGGTAGTCC |

**Table S5. Sequences and the copy numbers of standard DNAs of eukaryotic 18S rRNA**

| Primer name | Sequence |
|---|---|
| STD_euk1<br>10,000 copies/$\mu$l | GTACACACCGCCCGTCGCTCCTACCGATTGTGTGTGCGGGTGAAGGGACCGGATAGGTACT<br>TGGGGTTGTTTCCCAAGCTATTAACTAGAGAACTTTACTAAACCAGCACACATAGAGGAAG<br>GTGAAGTCGTAACAAGGTTTCCGTAGGTGAACCTGCAGAAGGATCA |
| STD_euk2<br>5,000 copies/$\mu$l | GTACACACCGCCCGTCGCTCCTACCGATTGTGTGTGCGGGTGAAGCGGCAGGATAGGCCTC<br>GAGGCTCGATTCCCAGATATGTCGACAGAGAACTTATCTAAACCGACACAAGTAGAGGAAG<br>GTGAAGTCGTAACAAGGTTTCCGTAGGTGAACCTGCAGAAGGATCA |
| STD_euk3<br>2,500 copies/$\mu$l | GTACACACCGCCCGTCGCTCCTACCGATTGCTTGTGCCGGTGAAGAGGATGGATGGGTGAG<br>CAGGGGATATTCCCCAGGGCAGAAGGCGAGAACTTTCGTAAACCGTCGCATATAGAGGAAG<br>GTGAAGTCGTAACAAGGTTTCCGTAGGTGAACCTGCAGAAGGATCA |
| STD_euk4<br>1,250 copies/$\mu$l | GTACACACCGCCCGTCGCTCCTACCGATTGCCTGTGCGGGTGAAGAGGATGGATCGGCACT<br>ACGGTGTTGTTCCCGTTTGGCCGGCGGGAGAATTTAAGTAAACCAGCGCAATTAGAGGAAG<br>GTGAAGTCGTAACAAGGTTTCCGTAGGTGAACCTGCAGAAGGATCA |
| STD_euk5<br>250 copies/$\mu$l | GTACACACCGCCCGTCGCTCCTACCGATTGTGTGTGCGGGTGAAGGGAAAGGATGGGGACT<br>GCGGCGCTTTTCCCGGCAACACTTAAAGAGAAATTCACTAAACCTACACAATTAGAGGAAG<br>GTGAAGTCGTAACAAGGTTTCCGTAGGTGAACCTGCAGAAGGATCA |

**Table S6. Sequences and the copy numbers of standard DNAs of fungal ITS**

| Primer name | Sequence |
|---|---|
| STD_asco1<br>20 copies/$\mu$l | CTCGGTCATTTAGAGGAAGTAAAAGTCGTAACAAGGTCTCCGTAGGTGAACCTGCGGAGG<br>GATCATTACCGAGCGAGGGACGGAGATGAAGCCTTGACACTTTGTGTCCGACACGGTTTG<br>CTTCGGGGGCGATTCTGCCGCAAAAGTTGCATTCCCCAAGATATTCGTCAAAACACTGCA<br>TCAACACGTCGGAACTAACTGTTAATGTTTCAAAACTTTCAACAACGGATCTCTTGGTTC<br>TGGCATCGATGAAGAACGCAGCGAA |
| STD_asco2<br>40 copies/$\mu$l | CTCGGTCATTTAGAGGAAGTAAAAGTCGTAACAAGGTCTCCGTAGGTGAACCTGCGGAGG<br>GATCATTACCGAGCGAGGGAAGTCCCCTACTACGTCAGTCTTTGTGTCCTAGCAAAGTTG<br>CTTCGGGGGCGACAATGCCGTACTCTCCGCATTCCCCTCATGCACGATCAAAACACTGCA<br>CATAGACGTCGGGGTAATCCGGTAATTCTCGAAAACTTTCAACAACGGATCTCTTGGTTC<br>TGGCATCGATGAAGAACGCAGCGAA |
| STD_asco3<br>100 copies/$\mu$l | CTCGGTCATTTAGAGGAAGTAAAAGTCGTAACAAGGTCTCCGTAGGTGAACCTGCGGAGG<br>GATCATTACCGAGCGAGGGAACTTCTGTCCGCGGACCTCCTTTGTGATCCGAAGAGGTTG<br>CTTCGGGGGCGATTTTGCCGGGGTTGACGCATTCCCCAGATACTATCTCAAAACACTGCA<br>TGATGACGTCGGTAGAGTCCTAGAACGAAGTAAAACTTTCAACAACGGATCTCTTGGTTC<br>TGGCATCGATGAAGAACGCAGCGAA |
| STD_asco4<br>200 copies/$\mu$l | CTCGGTCATTTAGAGGAAGTAAAAGTCGTAACAAGGTCTCCGTAGGTGAACCTGCGGAGG<br>GATCATTACCGAGCGAGGGAGGTTGTCATGCGTCCCGAACTTTGTGAGTAGTTACTTTTG<br>CTTCGGGGGCGAGTATGCCGAGGCGGAGGCATTCCCCCTAGCCGGGGTCAAAACACTGCA<br>CCTTAACGTCGGGTGAACGACCAAAAGCACTAAAACTTTCAACAACGGATCTCTTGGTTC<br>TGGCATCGATGAAGAACGCAGCGAA |
| STD_asco5<br>400 copies/$\mu$l | CTCGGTCATTTAGAGGAAGTAAAAGTCGTAACAAGGTCTCCGTAGGTGAACCTGCGGAGG<br>GATCATTACCGAGCGAGGGAATTTAACGGAGTCTGGGGGCTTTGTGTCGGCTCCGGTTTG<br>CTTCGGGGGCGATGCTGCCGGCGTGACGGCATTCCCCAGGTATCGCGTCAAAACACTGCA<br>TATCGACGTCGGAGACCGTCAGTAAAGAGGCAAAACTTTCAACAACGGATCTCTTGGTTC<br>TGGCATCGATGAAGAACGCAGCGAA |

**Table S7. Sequences and the copy numbers of standard DNAs of animal mitochondorial COI**

| Primer name | Sequence |
|---|---|
| STD_mlCOI1<br>200 copies/$\mu$l | GGAACAGGATGAACAGTTTACCCTCCACTGTCAACCAGTATTGCTCACAGAGGAGCTTCC<br>GTCGATTTAGCTATTTTTTCTTTACATTTAGCAGGCATCTCATCGATTATAGGGGCAATT<br>AACTTTATTACAACCGTTATTAATATACGAATGTGATGCATAAACGCGGATCGAATACCT<br>TTATTTGTTTGATCAGTCGTAATTACCAGGCTGATTTTTCTATTATCCCTACCCGTGCTA<br>GCCAGAGTGATCTCTATGCTTTCGTCTGATCGTAACTTAAATACATCATTCTACGACCCT<br>GCTGGTGGAGGGGACCCGATTTTGTATCAGCACCTATTTTGATTTTTTGGTCACCCTGAA<br>GTTTA |
| STD_mlCOI2<br>100 copies/$\mu$l | GGAACAGGATGAACAGTTTACCCTCCACTCTCTTCACCAATTGCTCACCGCGGAGCATCA<br>GTGGATTTAGCCATTTTTTCGTTTCATTTAGCAGGCATCTCCTCTATTTTAGGTGCATTA<br>AACTTTATTACAACCGTTATTAATATACGATGACCAACAATAGTTAGAGATCGAATACCT<br>TTATTTGTTTGATCAGTAGCTATTACCAGAGTCTTCTTTATATTATCACTACCAGTTCTA<br>GCTAGAGTAATCTCTATGCTTTCGTCTGATCGTAACTTAAATACATCATTCTACGACCCG<br>GCCGGGGGAGGAGACCCCATTTTCTATCAGCACCTATTTTGATTTTTTGGTCACCCTGAA<br>GTTTA |
| STD_mlCOI3<br>50 copies/$\mu$l | GGAACAGGATGAACAGTTTACCCTCCACTATCCCCATCTATTGCACACCATGGAGCCTCT<br>GTTGATTTAGCAATTTTTTCTTTACATTTAGCCGGATTGTCCTCTATTGTAGGGGCTATA<br>AACTTTATTACAACCGTTATTAATATACGAATGTCCTCTATATAGGTAGATCGAATACCT<br>TTATTTGTCTGATCTGTGGTCATTACGAGAGTGTTTCTAATTCTATCCCTACCGGTGCTA<br>GCCAGAGTGATCTCTATGCTTTCGTCTGATCGTAACTTAAATACATCATTCTACGACCCT<br>GCGGGAGGGGGCGACCCAATTTTGTATCAGCACCTATTTTGATTTTTTGGTCACCCTGAA<br>GTTTA |
| STD_mlCOI4<br>25 copies/$\mu$l | GGAACAGGATGAACAGTTTACCCTCCACTTTCACCAAGTATTGCTCACGGGGGAGCCTCT<br>GTCGATTTAGCAATTTTTTCTCTTCATTTAGCTGGTATTTCATCAATTTTAGGCGCACTT<br>AACTTTATTACAACCGTTATTAATATACGAGCACCCGAGATATGCTGTGATCGAATACCT<br>TTATTTGTTTGATCTGTGGAGATTACCAGGGTTATCCTACTTCTATCGCTACCCGTCCTA<br>GCTAGAGTAATCTCTATGCTTTCGTCTGATCGTAACTTAAATACATCATTCTACGACCCG<br>GCAGGTGGTGGGGACCCTATTTTTTATCAGCACCTATTTTGATTTTTTGGTCACCCTGAA<br>GTTTA |
| STD_mlCOI5<br>5 copies/$\mu$l | GGAACAGGATGAACAGTTTACCCTCCACTTTCTCCTATAATTGCTCACTGAGGAGCATCT<br>GTTGATTTAGCCATTTTTTCTCTACATTTAGCGGGTATATCATCTATTCTAGGAGCAGTA<br>AACTTTATTACAACCGTTATTAATATACGAACGGCCTTAATAGTTCCGGATCGAATACCT<br>TTATTTGTCTGATCAGTCGCCATTACGAGCGTCTTTCTTTTATTATCCCTACCCGTCCTA<br>GCGAGAGTAATCTCTATGCTTTCGTCTGATCGTAACTTAAATACATCATTCTACGACCCG<br>GCTGGTGGTGGTGACCCGATTTTGTATCAGCACCTATTTTGATTTTTTGGTCACCCTGAA<br>GTTTA |

#### 3. Summary of sequence quality

I performed MiSeq run four times to generate sequences from the four metabarcoding regions (i.e., 16S rRNA, 18S rRNA, ITS and COI). Summary of the sequence runs is provided below.

**Table S8. Summary of sequence quality**

| MiSeq run name | Cycles | Target region | Pass filter (%) | >Q30 (%) | Yield (Gbp) | Total reads | Reads PF |
|---|---|---|---|---|---|---|---|
| RMR-076 | 500 | 16S rRNA | 95.69 | 84.77 | 7.69 | 15,636,422 | 14,962,924 |
| CMR-002 | 300 | 18S rRNA | 89.64 | 94.86 | 5.67 | 20,158,344 | 18,070,998 |
| RMR-078 | 600 | ITS | 95.13 | 79.73 | 16.08 | 27,533,984 | 26,191,796 |
| RMR-099 | 500 | COI | 90.07 | 89.62 | 9.45 | 20,410,736 | 18,383,068 |
| Total | | | | | 38.89 | 83,739,486 | 77,607,998 |

Sequence reads from standard DNAs and non-standard DNAs are provided below.

**Table S9. Sequence reads from standard DNAs and non-standard DNAs**

| MiSeq run name | Standard DNAs (all libs) | Non-standard DNAs (all libs) | Standard DNAs (sample only) | Non-standard DNAs (sample only) | Standard/ Total |
|---|---|---|---|---|---|
| RMR-076 | 3,259,980 | 4,750,032 | 2,516,936 | 4,718,786 | 35% |
| CMR-002 | 5,391,480 | 7,437,546 | 4,485,282 | 7,394,159 | 38% |
| RMR-078 | 6,180,704 | 7,732,723 | 5,186,395 | 7,710,798 | 40% |
| RMR-099 | 3,343,130 | 7,693,367 | 2,759,167 | 7,472,133 | 27% |
| Per sample | 26,418 | 40,136 | 24,305 | 44,384 | 35% |

"all libs" indicates that sequence reads of all libraries including true samples, negative and positive controls are included. "sample only" indicates that sequence reads of only true samples are included (see sample metadata provided in https://github.com/ong8181/interaction-capacity/SpplementaryData).

#### 4. Rarefaction curve

Rarefying sequence reads is a common approach to evaluate microbial diversity. I examined rarefaction curves and found that the sequencing captured most of the prokaryotic diversity (Fig. S2). Also, a previous study suggested that simply rarefying microbial sequence data is inadmissible (McMurdie & Holmes 2014). Another important point is that my analyses involve conversion of sequence reads to DNA copy numbers and thus are different from other commonly used analyses in microbiome studies. Considering these conditions and the previous study, sequence reads were subjected to most of the downstream analysis in the present study without performing further corrections using rarefaction or other approaches.

#### 5. Negative control results

Prior to the library preparation, work-spaces and equipment were sterilized. Filtered pipet tips were used, and separation of pre- and post-PCR samples was carried out to safeguard against cross-contamination. To monitor cross-contamination during field sampling and library preparation process, 10 PCR-level negative controls without internal standard DNAs, 22 PCR-level negative controls with internal standard DNAs, and 30 field-level negative controls were employed for each MiSeq run. PCR-level negative controls were made by adding MilliQ water instead of extracted DNA. For field-level negative controls, a 500-ml plastic bottle containing MilliQ water was carried to the field with the other five 500-ml plastic bottles, and taken back to the laboratory after the water sampling. Then, the MilliQ water in the bottle were filtered using the cartridge filter ($\phi$ 0.22-$\mu$m and $\phi$ 0.45-$\mu$m). The filter cartridges were treated in an identical way with the other samples during the DNA extraction and library preparation processes. Field-level negative controls were collected consecutively during the first 15 sampling events (from 24 May 2017 to 6 June 2017, and the first pre-monitoring sampling on 22 May 2017). After the first 15 sampling events, field-level negative controls were collected once in a week (i.e., on Monday).

All of the negative controls generated negligible sequences compared with the field samples and positive controls for the four MiSeq runs (Fig. S2), suggesting that there was no serious cross-contamination during the water sampling, DNA extraction and library preparation processes.

#### 6. Positive control results

Positive control samples were taken to monitor for possible degradation during the sample storage. Water samples were collected from Plot 2 on 23 May 2017 before planting rice seedlings. Ten subsamples were separately filtered using the cartridge filter ($\phi$ 0.22-$\mu$m and $\phi$ 0.45-$\mu$m), and stored at -20°C in the same way as the other samples. DNAs of the positive controls were extracted subsequently and analyzed in the same way as the other samples. DNA extraction dates for the positive controls were as follows: 23 May 2017, 27 June 2017, 12 July 2017, 9 August 2017, 22 August 2017, 11 September 2017, 29 September 2017, 11 October 2017, 25 October 2017, and 8 November 2017 (i.e., from an immediate DNA extraction to 5.5 months storage at $-20^{□}$C).

The 10 most abundant ASVs were extracted and their abundances were visualized in Fig. S2I-L. The most abundant ASVs were robustly detected from most of the positive samples and there were no dramatic reductions in the DNA copy numbers if DNAs were extracted before 3 months after water sampling (red points in the figures). DNAs of the other samples were extracted within 1-2 months after water sampling in the present study, suggesting that there is no serious DNA degradation during the sample storage.

#### 7. Total DNA quantifications

Total DNAs were quantified using a Quant-iT assay kit (Promega, Madison, Wisconsin, USA). Three $\mu$l of each extracted DNA (from $\phi$ 0.22-$\mu$m Sterivex) was mixed with the fluorescent reagent (fluorescent reagent:buffer solution = 1:399) and DNA concentration was measured following the manufacturer's protocol.

The results were compared with the total DNA copy numbers estimated by the quantitative MiSeq sequencing (qMiSeq) of the four marker regions (i.e., 16S, 18S, ITS and COI). An assumption behind the analysis is that most cellular organisms were captured by sequencing the four marker regions. Fig. S3C shows the correlation between the total DNA concentrations and total DNA copy numbers. The total DNA copy numbers estimated by qMiSeq were correlated with the total DNA concentrations very well, showing the quantitative capacity of qMiSeq.

#### 8. Quantitative PCR (qPCR)

qPCR of the 16S region was performed using the same primer set with qMiSeq (515F-806R primers) (Bates *et al.* 2011; Caporaso *et al.* 2011). Two $\mu$l of each extracted DNA (from $\phi$ 0.22-$\mu$m Sterivex) was added to an 8-$\mu$l qPCR reaction containing 1 $\mu$l of 5 $\mu$M 515F primer, 1 $\mu$l of 5 $\mu$M 806R primer, 5 $\mu$l of Platinum SuperFi PCR Master Mix (ThermoFisher Scientific, Waltham, Massachusetts, USA), 0.5 $\mu$l of 20 $\times$ EvaGreen (Biotium, San Francisco, California, USA), and 0.5 $\mu$l of MilliQ water. The thermal cycle profile after an initial 30 s denaturation at 98°C was as follows (60 cycles): denaturation at 98°C for 10 s; annealing at 60°C for 10 s; and extension at 72°C for 15 s (fluorescent measured at this step). The measurements were performed in duplicate to mitigate technical errors. The standard dilution series were prepared using STD_pro1 used in qMiSeq.

Fig. S3D shows the relationship between the total 16S copy numbers estimated by qPCR and qMiSeq. Similarly to Fig. S3C, Fig. S3D shows that there is a highly significant positive correlation between the results of qPCR and qMiSeq. The absolute values estimated by qPCR were generally higher than those estimated by qMiSeq. This could be due to non-target amplifications during the qPCR: although 515F-806R primers predominantly amplify prokaryote 16S, the quantification by EvaGreen qPCR may include non-target DNA. A similar trend was also found in the previous study that used a universal fish primer set (Ushio *et al.* 2018). On the other hand, during the library preparation for qMiSeq, only the targeted amplicons were excised using E-Gel SizeSelect. Thus, non-target amplicons which might be included in qPCR quantifications were not included in qMiSeq quantifications. This could be a cause of the difference in the estimated DNA copy numbers between qPCR and qMiSeq. Nonetheless, both methods in general showed consistent results, again showing the quantitative capacity of qMiSeq.

#### 9. Shotgun metagenomic analysis

Shotgun metagenomic analysis was performed for a subset of the samples to check whether and how PCR-based assessments of community composition biased the results. Only four samples, of which diversity was high, were analyzed because much deeper sequencing is necessary for the shotgun metagenomic analysis (e.g., $>×$100 deeper sequencing). Briefly, approximately 10–30 ng of total DNA were used as inputs, and Illumina DNA Prep (Illumina, San Diego, CA, USA) was used to prepare the library for the shotgun metagenome. The library was prepared by following the manufacturer's protocol. The double-stranded DNA concentration of the library was then adjusted to 4 nM and the DNA was sequenced on the MiSeq using a MiSeq V2 Reagent kit for 2 × 250 bp PE (Illumina, San Diego, CA, USA).

In total, 20,601,323 reads (10.3 Gb for 4 samples) were generated (Table S10) and the quality of sequencing was generally high (>Q30 = 80.4%). The low quality reads and adapter sequences were removed using fastp (https://github.com/OpenGene/fastp) (Chen *et al.* 2018). After the quality filtering, 14,315,030 sequence reads ramained and the filtered sequences were analyzed using phyloFlash (http://hrgv.github.io/phyloFlash/) (Gruber-Vodicka *et al.* 2020). phyloFlash extracted and assembled 16S sequences from the sequence reads. Then, taxonomy and the number of sequence reads assigned to each taxa were summarized in the same pipeline.

**Table S10. Sequence results of the shotgun metagenomic analysis**

| Sample ID | Sampling date | Plot | Sequence reads | Data size |
|---|---|---|---|---|
| S115 | 2017/7/14 | 1 | 5,197,043 | 2.6 Gb |
| S116 | 2017/7/15 | 1 | 4,993,487 | 2.5 Gb |
| S565 | 2017/7/14 | 5 | 4,764,166 | 2.4 Gb |
| S566 | 2017/7/15 | 5 | 5,646,627 | 2.8 Gb |
| total | | | 20,601,323 | 10.3 Gb |

Fig. 3F-I show the comparison between qMiSeq and the shotgun metagenomic analysis. In summary, the numbers of detected taxa were comparable between the two methods: 82-136 by qMiSeq and 99-118 by the shotgun metagenomic analysis. In addition, the community compositions detected by the two methods were also similar. For both methods, Proteobacteria and Bacteroidetes were the two most dominant phyla. Although qMiSeq detected Verrucomicrobia slightly more frequently than the shotgun metagenomic analysis did, the difference was not large. Thus, the shotgun metagenomic analysis also showed that qMiSeq reasonably captured the diversity and composition of the ecological communities in the present study.

#### 10. Software packages

Descriptions of main software packages used in this study. Detailed information for the packages and analysis environments are available in Github (https://github.com/ong8181/interaction-capacity).

##### Sequence processing

- bcl2fastq (bcl2fastq2-v2-18-0-12-linux-x86-64)
- Claident v0.2.2019.05.10 (https://www.claident.org/)
- cutadapt v2.5
- dada2 v1.11.5
- phyloseq v1.28.0

##### Time series analysis

- rEDM v0.7.5

**General**

- tidyverse v1.3.0
- RCpp v1.0.3
- pforeach v1.3
- glmnet v3.0.1

#### 11. SI Dataset: Legend for Supporting Dataset (Dataset S1)

Sample metadata is provided as a separate CSV file (see SI_Dataset.csv in https://github.com/ong8181/interaction-capacity/tree/master/SupplementaryData). The metadata include sampling date, filtered water volumes for each filter type, sequence reads, and DADA2 processing summary.

#### 12. SI Movie: Legend for Supporting Movie (Movie S1)

Time-varying interaction network of Plot 1. Blue and red arrow indicates positive and negative causal interactions, respectively.
